# Cell-type-resolved transcriptional reprogramming in resistant soybean roots reveals cambial activation and early syncytium initiation upon nematode infection

**DOI:** 10.64898/2026.04.08.717279

**Authors:** Vikas Devkar, Leonidas D’Agostino, Sushil Satish Chhapekar, Jiamei Li, Moises Frausto, Lenin Yong□Villalobos, Luis Herrera-Estrella, Fiona L. Goggin, Henry Nguyen, Gunvant B. Patil

## Abstract

Soybean cyst nematode (SCN) is the most destructive pathogen of soybean, yet the cellular basis of host resistance remains poorly understood. Here, we present a high-quality, cell-type–resolved atlas of root responses during early SCN infection in the highly resistant genotype PI437654, capturing transcriptional states across all major tissues, including rare syncytial cells. Our analyses reveal that resistance is mediated not by a localized defense but by coordinated, multicell reprogramming spanning invasion layers, vascular tissues, and feeding site–associated cells. We identify the vascular cambium as the primary cellular origin of SCN-induced syncytia, resolving a long-standing question in nematology. Mechanistically, resistance arises from disruption of key processes required for feeding site establishment, secretory stress via imbalanced vesicle trafficking, suppression of endoreduplication to prevent hypertrophic syncytial growth, and activation of autophagy to maintain cellular homeostasis. Spatially organized hormone signaling networks, including jasmonic acid, salicylic acid, and ethylene pathways, further reinforce defense, with GmJAZ1 functioning as a central regulator of JA–SA crosstalk. Collectively, PI437654 enforces resistance by targeting host cell identity, nutrient sink formation, and sustained parasitism, deploying a multilayered, tissue-specific defense strategy. This study provides a mechanistic, systems-level framework for SCN resistance and establishes a single-cell resource capturing rare root cell states, offering actionable targets for engineering durable nematode resistance.

**Key points:**

- Soybean cyst nematode (SCN) is the most destructive pathogen of soybean worldwide, yet the cellular basis of early host responses and feeding site initiation remains poorly understood.
- Using single-nucleus RNA sequencing (snRNA-seq), we generated a cell-type–resolved atlas of early SCN infection in roots of a unique and highly resistant soybean genotype PI437654. Trajectory analysis integrated with syncytium marker genes revealed that cambium cells are selectively targeted as the cellular origin of syncytium formation.
- SCN infection triggers extensive cell-type–specific transcriptional reprogramming, particularly in vascular tissues (xylem, phloem, and cambium), involving pathways related to cell cycle and endoreduplication, vesicle trafficking, autophagy, and phytohormone signaling.
- Functional validation demonstrated enhanced autophagy activation in infected roots via increased GFP-GmATG8a–labeled autophagic puncta, while overexpression of the jasmonic acid regulator GmJAZ1 significantly enhanced SCN resistance in susceptible soybean.
- Together, these findings define the cellular origin of SCN-induced syncytia and reveal coordinated cell-type-specific defense programs, providing a mechanistic framework for engineering durable resistance to SCN.

## Introduction

Soybeans rank second in production after corn in the USA, serving as a vital source of protein, livestock feed, as well as edible and biofuel oil (Anderson *et al*., 2019). Despite its agronomical and economical significance, soybean yield is adversely affected by biotic and abiotic stresses, emphasizing the need for targeted trait improvement to withstand yield and quality (Deshmukh *et al*., 2014; Bandara *et al*., 2020; Liu *et al*., 2020). Among the biotic stresses, soybean cyst nematode (SCN) (*Heterodera glycines*) is the most destructive soybean pathogen in the USA, causing approximately $1.5 billion in yield losses annually (Tylka & Marett, 2025). County-level surveys show that SCN is widespread across North America, and its presence has been confirmed in the soil of nearly all soybean growing counties of the United States (Tylka & Marett, 2025).

Present SCN management practices rely primarily on the use of resistant soybean cultivars and crop rotation with nonhost crops (Patil *et al*., 2019b; Arjoune *et al*., 2022; Thapa *et al*., 2022). However, most SCN-resistant soybean cultivars in the US are derived predominantly from the resistance sources PI 88788 and, to a lesser extent, Peking, which have been extensively utilized in soybean breeding programs (Tylka & Mullaney, 2015). These resistance sources harbor the major loci *rhg1* and *Rhg4*, whose genetic mechanisms of SCN resistance have been well characterized (Cook *et al*., 2012; Liu *et al*., 2012). Nevertheless, decades of widespread reliance on PI 88788-derived resistance have imposed strong selection pressure on SCN populations, leading to the emergence of virulent nematode populations capable of reproducing on PI 88788-based cultivars (Niblack *et al*., 2008; McCarville *et al*., 2017). Therefore, identifying alternative sources of resistance becomes essential to counter nematode virulence. Among the limited number of highly resistant soybean genotypes, PI437654 represents one of the most effective resistance sources, conferring broad-spectrum resistance against multiple SCN populations and HG types (Mahalingam & Skorupska, 1996; Patil *et al*., 2019a). Surveys of global soybean germplasm indicate that PI437654 contains unique allelic variants and an expanded Rhg4 copy number, including four copies of the associated QTL (Patil *et al*., 2019b). Thus, given the reduced effectiveness of PI88788-derived resistance and the emergence of PI437654 as one of the most effective sources of SCN resistance, unraveling the cell-type-specific molecular mechanisms governing SCN resistance, particularly during early infection and attempted feeding site establishment, is essential for developing durable resistance strategies.

SCN is an obligate biotrophic, plant-parasitic roundworm that establishes a specialized feeding structure, known as a syncytium, within the host root vasculature (Mitchum *et al*., 2013). This feeding site is formed through extensive remodeling and fusion of host cells and serves as a long-term nutrient source that sustains nematode development. Syncytium formation is initiated by nematode-secreted effector proteins delivered into host root cells, which reprogram host transcriptional networks and manipulate cellular processes including cell cycle regulation, vesicle trafficking, and metabolism to establish and maintain the feeding site (Hewezi & Baum, 2013; Kyndt *et al*., 2013). Importantly, cytological analyses of the highly resistant genotype PI 437654 demonstrate that the initial stages of syncytium establishment are largely conserved between resistant and susceptible interactions, with both genotypes exhibiting cell wall dissolution, hypertrophy of selected cells, and localized hyperplasia of surrounding tissues during early infection (Mahalingam & Skorupska, 1996; Kim, Kim & Riggs, 2010). These findings indicate that resistance does not prevent nematode entry or the initiation of feeding site formation. Rather, divergence occurs shortly after establishment: in PI 437654, syncytium expansion is rapidly restricted, accompanied by early onset of defense responses including necrosis, wall appositions, nuclear disintegration, and cytoplasmic collapse, ultimately leading to premature degeneration of the feeding site within days of infection. In contrast, susceptible genotypes support continued syncytium enlargement and metabolic activation, enabling successful nematode development (Mahalingam & Skorupska, 1996; Kim, Kim & Riggs, 2010).

Understanding how these host cellular processes are coordinated during early infection and how initially similar developmental programs diverge toward either functional or abortive syncytia is essential for deciphering mechanisms of SCN resistance. Previous studies have characterized SCN-induced transcriptional responses using bulk root transcriptomics (Zhang *et al*., 2017; Torabi *et al*., 2023; Nissan *et al*., 2025) and laser-capture microdissection (LCM)–based syncytium transcriptome profiling (Klink *et al*., 2010). While these approaches have provided valuable insights into host gene expression during infection, they lack the spatial and cellular resolution required to define the precise cellular origin of syncytium initiation and to resolve cell-type-specific host responses within complex root tissues. Recent advances in single-cell and single-nucleus transcriptomics now enable high-resolution dissection of plant–microbe interactions across individual cell types and tissues, uncovering previously hidden regulatory networks governing host responses to pathogens (Tang *et al*., 2023; Devkar *et al*., 2025a; Nobori *et al*., 2025).

Here, we applied single-nucleus RNA sequencing (snRNA-seq) to characterize cellular and transcriptional heterogeneity in SCN-uninfected (SCN−) and SCN-infected (SCN+) roots of the highly resistant soybean genotype PI437654. Integrating snRNA-seq with trajectory analysis, we identify cambium cells as the primary cellular origin of SCN infection and syncytium initiation. Our analyses further reveal extensive cell-type-specific transcriptional reprogramming during infection, particularly in vascular tissues, involving pathways associated with vesicle trafficking (endocytosis and exocytosis), cell cycle progression, endoreduplication, autophagy, and hormone biosynthesis and signaling. Functional validation confirmed activation of the autophagy pathway, visualized through increased GFP-GmATG8a–labeled autophagic puncta in SCN-infected roots. In addition, we demonstrate that manipulation of JA signaling through the constitutive expression of the master regulator *GmJAZ1* enhances resistance to SCN. Collectively, our study provides an in-depth cell-type–resolved framework of SCN infection in soybean roots, revealing the cellular origin of syncytium formation and identifying key regulatory pathways underlying host defense. These findings establish a mechanistic foundation for leveraging cellular-scale insights to develop next-generation soybean cultivars with durable resistance to SCN.

## Results and Discussion

### snRNA-seq analysis of soybean root cells during SCN infection

To validate the resistance phenotype used for downstream analyses, we re-evaluated response of PI437654 to SCN. As expected, PI437654 exhibited strong resistance, with a markedly reduced penetration of juveniles (J2/J3) in roots (Fig. 1), followed by a reduced number of female cysts relative to susceptible controls. These results confirm the robust resistance of PI437654, and the SCN-infected (SCN+) and non-infected (SCN-) roots from PI437654 (5 days post infection) were used to investigate early cellular responses to SCN infection using snRNA-seq. After quality filtering, a total of 6,912 high-quality nuclei were retained for downstream analysis. Alignment of sequencing reads to the soybean reference genome detected expression of 45,674 genes, representing approximately 82% of all protein-coding genes in soybean. On average, each nucleus contained a median of 351 unique molecular identifiers (UMIs) and 565 expressed genes (**Table S1**). Dimensionality reduction using uniform manifold approximation and projection (UMAP) resolved the nuclei into 17 transcriptionally distinct clusters, including syncytium cells derived from cambium cells (**Figs 1b, S1; syncytium cells highlighted with a dotted circle)**. Cluster identities were assigned using established cell-type marker genes from recently published soybean single-cell and single-nucleus transcriptomic datasets (**Fig. 1c; Table S2**) (Liu *et al*., 2023; Sun *et al*., 2023; Cervantes-Pérez *et al*., 2024; Zhang *et al*., 2025). These clusters corresponded to major root cell types, including epidermis (clusters 1–5), trichoblast/root hair cells (cluster 10), cortex (clusters 2 and 13), endodermis (clusters 6, 8, 9 and 14), cambium (clusters 7 and 12), pericycle (clusters 11 and 15), phloem (cluster 16), and xylem (cluster 17) (**Figs 1 and S1-S2**). A graphical representation of the root cross-section under infection from SCN is detailed in **Fig. 1d**, with the feeding zone of the SCN, the cambium, highlighted. Analysis of SCN-responsive marker genes revealed pronounced transcriptional activation across multiple root cell types following infection (Fig. 1e). Notably, the SCN+ samples exhibited broad, multi-cell-type transcriptional activation, suggesting that resistance involves coordinated responses across several root tissues rather than a localized defense program **(Fig 1b, e)**.

**Figure 1.**
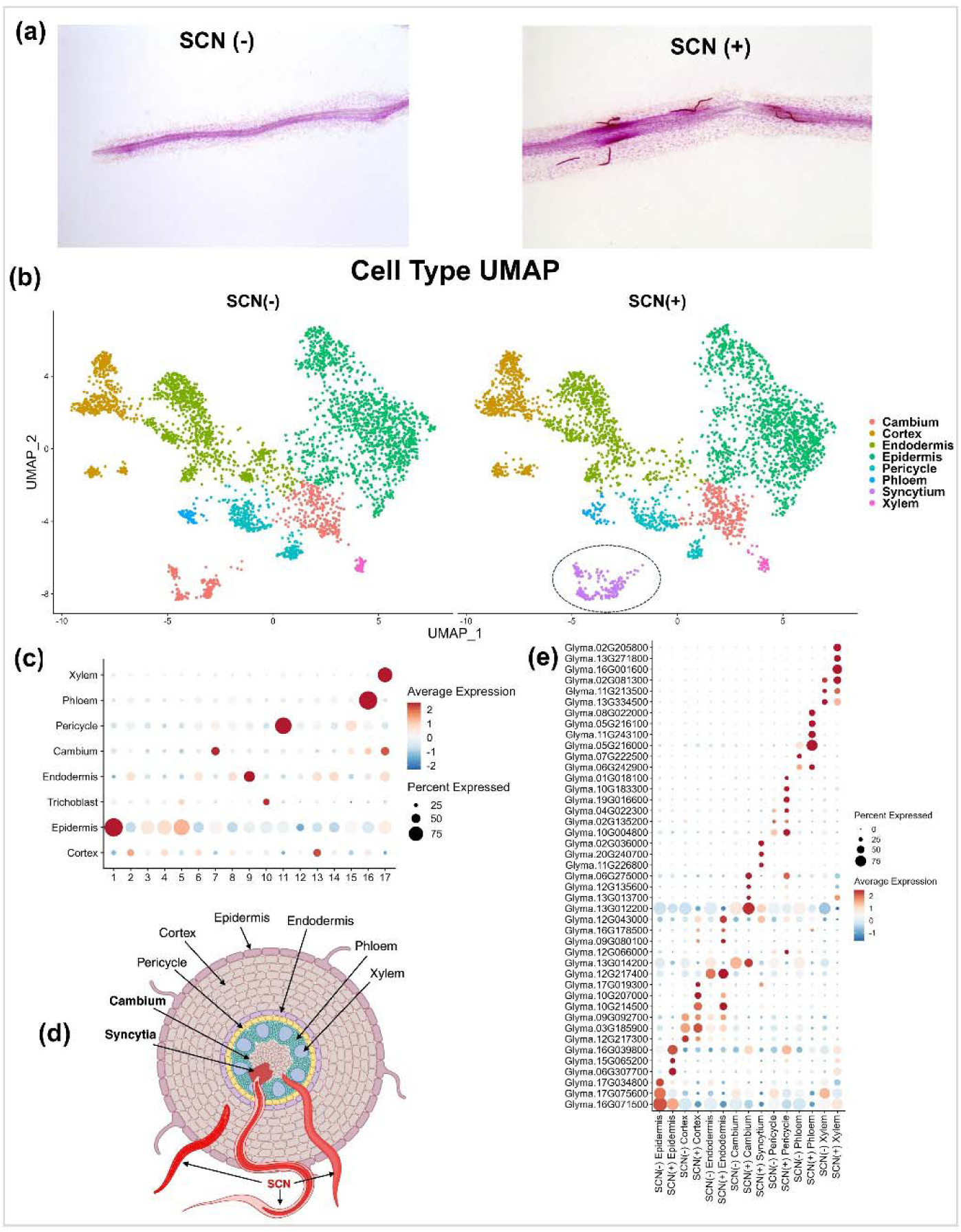
Single nucleus RNA-sequencing (SnRNA-seq) profiles of soybean roots infected with soybean cyst nematode (SCN) (+) and non-infected SCN (-) in resistant cultivar PI437654. (a) Soybean cyst nematode (SCN) infected SCN (+) for 5 days and non-infected SCN (-) roots stained with fuchsin dye. (b) UMAP representation of SCN (-) and SCN (+) soybean root cell type clusters. Clusters of diverse colors denote cell identities. (c) Dot plot represents the combined expression of marker genes in various clusters specific to cell-types. The size of the dots indicates the percentage of cells in each cluster that express the genes. The colors show the relative gene expression. (blue = lower expression, Red = higher expression). (d) Cartoon exhibits different cell types in soybean root and migration and infection site of SCN. (e) Dot plot shows top three marker genes specifically expressed in specific cell-type in soybean roots treated with SCN (+) and SCN (-).

To further characterize the biological processes underlying these transcriptional changes, we performed Gene Ontology (GO) enrichment analysis at single-nucleus resolution. SCN infection induced extensive cell-type-specific transcriptional reprogramming, with the strongest responses observed in vascular and pericycle tissues. In vascular and pericycle clusters, GO terms associated with clathrin-dependent endocytosis, receptor-mediated endocytosis, vesicle uncoating, and membrane disassembly were strongly enriched (**Fig. S3**). These pathways are consistent with the extensive membrane trafficking required for nematode effector uptake and syncytium establishment (Gheysen & Mitchum, 2011; Goverse & Smant, 2014). Phloem cells showed enrichment for conserved GO categories related to sucrose transport and phloem loading, indicating metabolic reprogramming that may facilitate nutrient allocation toward the developing feeding site. In contrast, cambium and vascular cell populations displayed enrichment of ribosome biogenesis, rRNA processing, and translational elongation, suggesting enhanced biosynthetic activity associated with syncytial development **(Fig. S3)**. Collectively, these findings indicate that SCN infection induces coordinated, cell-type-specific remodeling of vesicle trafficking, metabolic allocation, and stress-associated signaling pathways across root tissues, thereby creating a cellular environment conducive to feeding site establishment.

### Cambium-derived syncytia cells

Formation of a syncytium feeding site is a critical step for the successful establishment of SCN infection (Sobczak & Golinowski, 2011). To determine the cellular origin of syncytium formation upon infection, we analyzed our snRNA-seq dataset by considering previously reported syncytium-specific marker genes derived from LCM-based transcriptomic studies (Klink *et al*., 2005; Klink *et al*., 2007; Klink *et al*., 2010) (**Fig. S4; Table S3**). These earlier studies isolated SCN-induced syncytial tissues from soybean roots and identified genes specifically enriched in feeding sites relative to surrounding root tissues. While classical histological studies have suggested that syncytia originate within vascular tissues (Gheysen & Mitchum, 2011; Kyndt *et al*., 2013), the precise cellular identity of the initiating host cell population has remained unresolved. To answer this question, we focused on vascular-associated cell populations and performed re-clustering of vascular cell types within our snRNA-seq dataset (**Fig. 2a**). Expression profiling of 217 previously defined syncytium marker genes revealed a striking enrichment within SCN-infected cambium cluster 12, whereas other vascular cell types, including xylem, phloem, and pericycle, showed comparatively weaker expression of these markers **(Fig. 2b)**. Based on this strong transcriptional signature, we designated SCN-infected cambium cluster 12 as syncytial cells, indicating that cambium cells represent the most probable cellular origin of syncytium formation in soybean roots.

**Figure 2.**
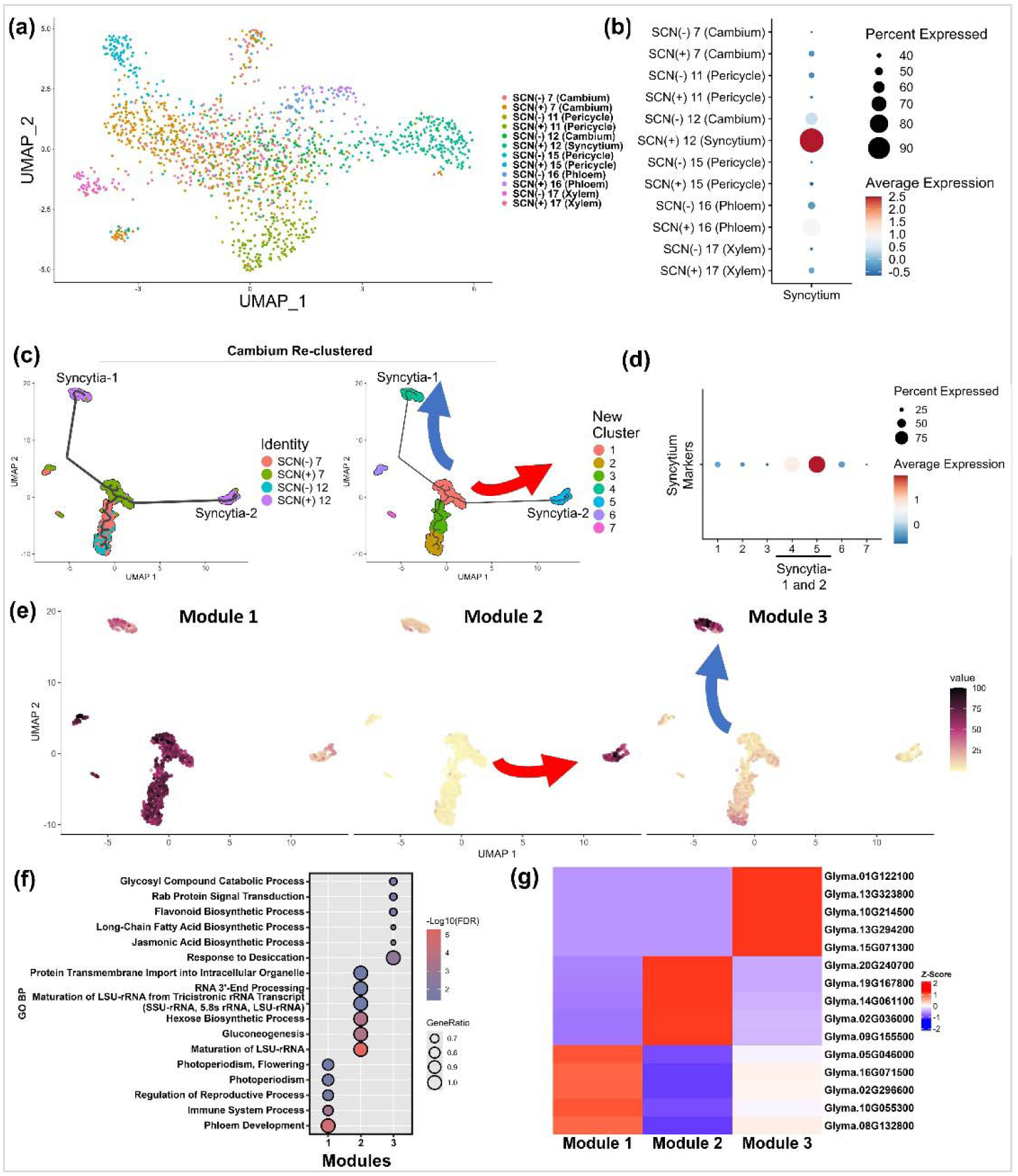
Analysis of cambium derived syncytia cells in SCN-resistant reaction (PI437654). **(a)** UMAP plot shows re-clustering of vascular cells in soybean roots treated with SCN (+) and SCN (-). **(b)** Dot plot shows expression of syncytia specific SCN marker genes from Klinck et al. 2005; 2007; 2010 in different vascular cell-types in soybean roots treated with SCN (+) and SCN (-). The colors show the relative gene expression. (blue = lower expression, Red = higher expression). **(c)** Trajectory analysis using a dynamical model shows that cambium cells subdivide into seven distinct subclusters and give rise to two separate syncytium cell clusters, Syncytia-1 and Syncytia-2. **(d)** Dot plot depicts expression of syncytia specific SCN marker genes from Klinck et al. 2005; 2007; 2010 in cambium sub-clusters. **(e)** UMAP plot shows re-clustering of cambium cells, which segregate into three distinct modules based on differential gene expression patterns; arrows indicate separate developmental streams. The colors depict the relative gene expression. (yellow = lower expression, Purple = higher expression). **(f)** Gene ontology (GO) enrichment analysis of three transcriptional modules derived from seven cambium subclusters under SCN (-) and SCN (+) conditions. **(g)** Heatmap showing scaled gene expression of the top five marker genes for each of the three modules, with red denoting high expression and blue indicating low expression.

To further resolve the transcriptional heterogeneity within the cambial population, we re-clustered cambium cells (clusters 7 and 12), which resulted in seven distinct subclusters **(Fig. 2c)**. Trajectory analysis revealed that syncytial cells segregate into two transcriptionally distinct populations, hereafter referred to as syncytia-1 (subcluster 4) and syncytia-2 (subcluster 5) (**Fig. 2d)**. Both populations expressed canonical syncytium markers, although expression was markedly higher in syncytia-2, suggesting that these cells likely correspond to actively infected feeding cells containing the nematode, whereas syncytia-1 cells may represent surrounding or transitional cells, including pre-infection, early remodeling, or post-infection states (Fig. 2d).

To characterize the molecular programs underlying these cellular states, we performed differential gene expression analysis along the cambium trajectory, which identified three distinct transcriptional modules with unique expression dynamics **(Fig. 2e)**. Module 1 was associated with baseline cambial cells and likely represents uninfected cambium identity. In contrast, Module 2, strongly enriched in syncytia-2 cells, showed significant overrepresentation of biological processes related to ribosome biogenesis, including large subunit rRNA maturation and RNA 3′-end processing, together with carbohydrate metabolic pathways such as hexose biosynthesis and gluconeogenesis (Fig. 2f). These enrichments suggest substantial upregulation of protein synthesis and carbon metabolism, consistent with the concept that nematode feeding sites function as highly active metabolic and biosynthetic hubs that supply nutrients to the developing parasite (Hofmann *et al*., 2009; Hofmann *et al*., 2010; Vijayapalani *et al*., 2018).

In contrast, Module 3 (enriched in syncytia-1), displayed strong activation of pathways associated with Rab-mediated vesicle trafficking, lipid metabolism, and secondary metabolic processes, including long-chain fatty acid and flavonoid biosynthesis. Notably, this module also showed enrichment of jasmonic acid biosynthesis pathways, highlighting activation of hormone-associated defense and metabolic signaling **(Fig. 2f)**. These observations are consistent with previous studies showing that cyst nematodes actively manipulate host hormone signaling networks and defense-associated metabolic pathways during feeding site formation (Chin, Behm & Mathesius, 2018; Gheysen & Mitchum, 2019). Representative marker genes from each module are shown in Fig. 2g. Importantly, these results reveal a biphasic transcriptional response within cambium cells during SCN infection. Cells directly associated with the feeding site exhibit transcriptional programs consistent with high metabolic activity and nutrient production, whereas adjacent or transitional cambium cells display vesicle trafficking and defense-associated regulatory programs. This *in silico* spatial and functional diversification suggests that SCN-induced remodeling extends beyond the primary feeding cell, involving neighboring cambial cells that may contribute to feeding site expansion, structural maintenance, or host defense modulation. In summary, the preferential targeting of cambium cells for syncytium initiation may reflect the unique developmental properties of vascular cambium. Cambium is a meristematic tissue composed of pluripotent stem cells that continuously generate xylem and phloem throughout plant growth (Miyashima *et al*., 2012). Unlike terminally differentiated root tissues, cambial cells maintain developmental plasticity, sustained metabolic activity, and the capacity to re-enter the cell cycle (Miyashima *et al*., 2012; Crang, Lyons-Sobaski & Wise, 2018). These features likely make cambium cells particularly susceptible to nematode-driven reprogramming and provide a permissive cellular environment for extensive transcriptional and structural remodeling required for syncytium formation. Collectively, the study suggests that cambium cells constitute the primary cellular origin of SCN-induced syncytia, revealing previously unrecognized heterogeneity within feeding site-associated cells and uncovering transcriptional programs that underpin the early stages of nematode parasitism.

### Cell-type-specific reprogramming of key cellular processes during SCN infection

Following the identification and characterization of syncytium-forming cells, we next investigated the broader cellular programs that were highly enriched with SCN infection. Syncytium formation is known to require extensive host cell reprogramming, including modulation of vesicle trafficking, cell-cycle regulation, metabolic activity, and phytohormone signaling pathways (Goverse & Bird, 2011; Gheysen & Mitchum, 2019). Previous studies have also reported that nematode infection induces endoreduplication, membrane remodeling, and enhanced metabolic activity to support the development of hypertrophic feeding cells that function as nutrient sources for the parasite (Breuer, Braidwood & Sugimoto, 2014; Kyndt, Fernandez & Gheysen, 2014). In parallel, cellular recycling and defense-associated pathways, including autophagy and hormone signaling, contribute to maintaining cellular homeostasis and regulating host responses during pathogen invasion (Üstün, Hafren & Hofius, 2017; Marshall & Vierstra, 2018). Guided by these established mechanisms, we examined key cell-type-specific dynamics enriched for vesicle trafficking, autophagy, cell-cycle regulation, and phytohormone signaling across the snRNA-seq atlas to understand how these pathways are coordinated during SCN infection.

### SCN promotes endo and exocytosis

Endocytosis plays an important role during plant-pathogen interactions by mediating remodeling of plasma membrane proteins, internalization of receptors, and recycling of vesicles to modify immune signaling and defense responses (Leborgne-Castel, Adam & Bouhidel, 2010; Beck *et al*., 2012; Fan *et al*., 2015; Zhang, Xing & Lin, 2019). In plants, clathrin-mediated endocytosis (CME) is initiated at the plasma membrane through cargo recognition by adaptor complexes, such as AP2, TPLATE, TWD40, and LOLITA (Gadeyne *et al*., 2014; Fan *et al*., 2015). These adaptors recruit clathrin triskelions composed of clathrin heavy (CHC) and light chains (CLC), forming the clathrin coat that drives vesicle budding (Chen, Irani & Friml, 2011). Vesicle scission is subsequently mediated by dynamin-related proteins (DRPs), releasing clathrin-coated vesicles (CCVs) from the plasma membrane (Ramachandran, 2011). Subsequently, Auxilin and the ATP-dependent chaperone HSC70 disassemble the clathrin coat, allowing vesicle recycling and downstream trafficking (Xing *et al*., 2010; Adamowski *et al*., 2018).

To determine whether SCN infection modulates CME in a cell-type-specific manner, we analyzed the expression of CME-associated genes (Sato *et al*., 2009) across the snRNA-seq dataset **(Fig. 3a, c; Table S4).** Core clathrin coat genes, *GmCHC* and *GmCLC*, were enriched in xylem and syncytium cells **(Fig. 3a, c)**, indicating active vesicle formation within vascular tissues and the nematode-induced feeding site. Similarly, adaptor complex genes (*GmAP2A*, *GmAP2B*, *GmAP2M*, *GmAP2S*) in cortex, pericycle, and xylem cells, as well as plant-specific adaptor components (*GmTPLATE*, *GmTWD40*) in xylem, phloem, and syncytia were enriched **(Fig. 3a,c)**. In addition, vesicle scission factors *GmDRP1* and *GmDRP2* (Bednarek & Backues, 2010) were upregulated in syncytium and xylem cells, suggesting active clathrin-coated vesicle (CCV) budding during infection.

**Figure 3.**
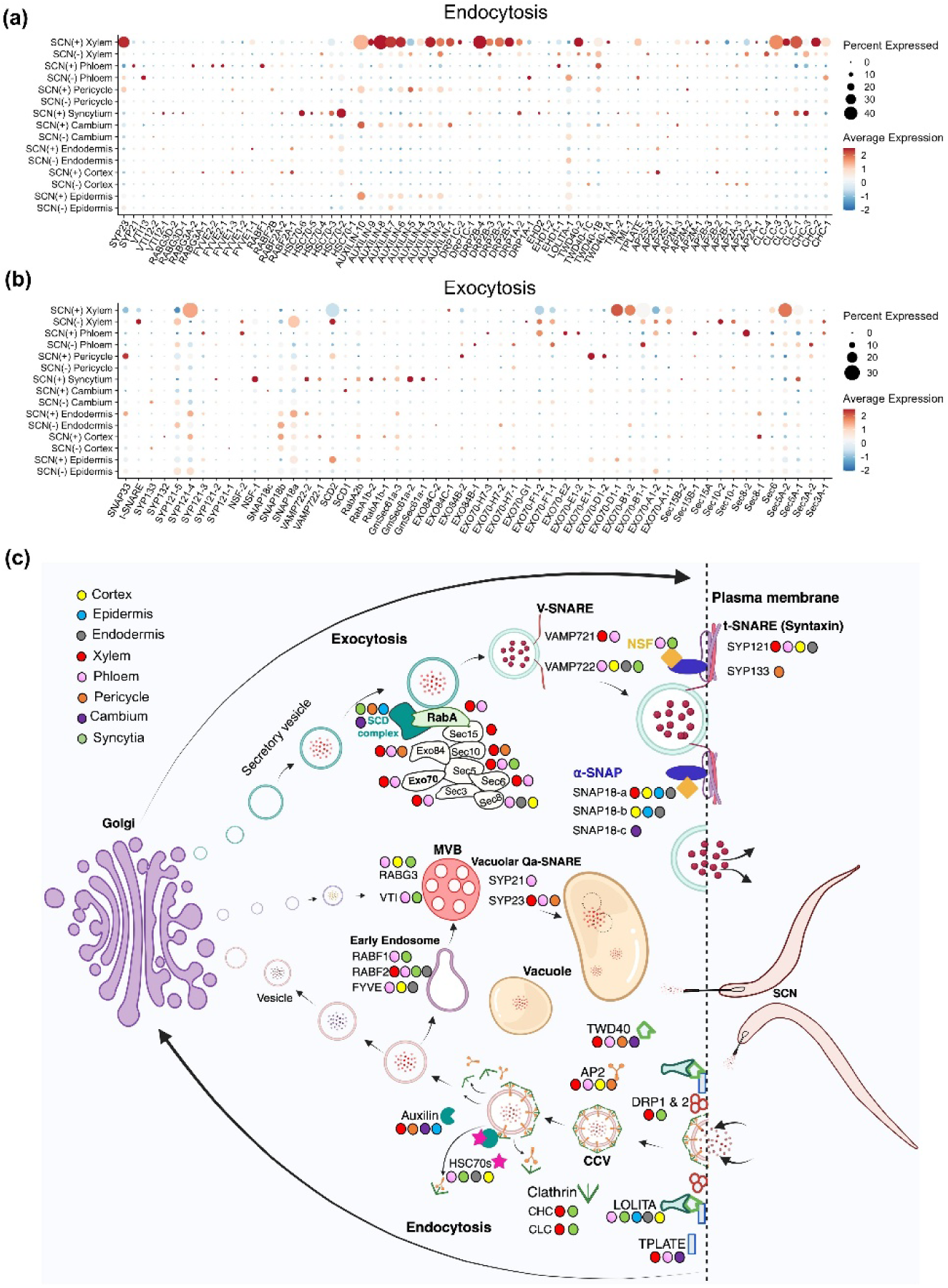
Perturbation of exocytosis and endocytosis vesicular trafficking pathways in different cell types after SCN infection. **(a)** A dot plot exhibits transcript abundance of clathrin mediated endocytosis (CME) and vacuolar trafficking pathway genes upon SCN (+) treatment. **(b)** Dot plot shows expression of exocytosis pathway genes upon SCN (+) treatment. The size of the dots depicts the percentage of nuclei in each cluster that express the gene. The colors indicate the relative expression. The Red dots denote higher expression while blue dots exhibit lower expression. **(c)** Schematic model illustrates coordinated regulation of clathrin-mediated endocytosis (CME), exocytosis, and vacuolar trafficking pathways in soybean cells after SCN infection (SCN+). Different colored circles indicate cell-type-specific expression of genes associated with these pathways, as identified from the SnRNA-seq analysis.

Consistent with elevated vesicle turnover, genes involved in CCV uncoating, GmAUXILIN, were differentially expressed in epidermis, xylem, pericycle, and cambium cells, whereas GmHSC70s, which cooperate with auxilin to disassemble clathrin coats, showed strong enrichment specifically in syncytium cells **(Fig. 3a)**. This pattern suggests rapid clathrin coat recycling and sustained CME activity within the nematode feeding structure. Following uncoating, vesicles are routed through the endosomal trafficking pathway for cargo sorting, recycling, or degradation. This process is coordinated by Rab GTPase genes, which regulate endosomal maturation toward multivesicular bodies (MVBs) and subsequent vacuolar trafficking, where membrane fusion is mediated by vacuolar Qa-SNAREs, including SYP21, SYP22, and SYP23 (Uemura et al., 2010; Viotti et al., 2010; Cui et al., 2014). These cell-type-specific expression patterns are consistent with previous studies describing activation of endosomal and vacuolar trafficking during plant–pathogen interactions (Leborgne-Castel, Adam & Bouhidel, 2010; Gu, Zavaliev & Dong, 2017), but our analysis provides single-cell resolution, revealing the specific vascular and syncytial cell populations in which these trafficking processes are most active. Together, these results suggest that SCN infection induces a coordinated activation of CME and endosomal trafficking pathways in vascular and syncytial cells. Mechanistically, enhanced CME in vascular cells may support syncytium formation and facilitate nutrient mobilization, internalization of immune receptors, and uptake or trafficking of nematode effectors. Such pathogen-driven manipulation of host vesicle trafficking has been widely observed during biotrophic interactions (Leborgne-Castel, Adam & Bouhidel, 2010; Beck et al., 2012; Fan et al., 2015), and our cell-type-resolved data indicate that this process is particularly intensified within the syncytium and adjacent vascular tissues that sustain the nematode feeding site **(Fig. 3C)**.

In plant cells, exocytosis is a central vesicle trafficking pathway that transports molecules (polysaccharides, membrane proteins, and defense cargoes) from the trans-Golgi network to the plasma membrane and apoplast (Uemura *et al*., 2019; Zhang, Xing & Lin, 2019). Secretory vesicles emerging from the Golgi are guided by RabA GTPases and the Stomatal Cytokinesis Defective (SCD) complex during post-Golgi trafficking (Mayers *et al*., 2017). Then vesicles are tethered by GTPase Secretory (*SEC* complex) and exocyst genes (EXO70 and EXO84) (Fendrych *et al*., 2013; Sharma *et al*., 2020). Membrane fusion is then mediated by vesicle-associated v-SNAREs VAMP721/722 interacting with plasma membrane t-SNAREs (SYP121, SYP133) (Kalde *et al*., 2007; El Kasmi *et al*., 2013). Following fusion, NSF and α-SNAP catalyze ATP-dependent disassembly of SNARE complexes to recycle t-SNAREs for subsequent rounds of secretion (Hanson *et al*., 1997).

Our analysis revealed pronounced cell-type-specific remodeling of the exocytosis machinery during SCN infection (Fig. 3b,c). Components associated with vesicle targeting and tethering were selectively induced in vascular tissues, including *GmSEC5A* and *GmSEC8*, while *GmSCD1* and several *GmRabA* isoforms were preferentially enriched in syncytial cells. In parallel, members of the *EXO70* family displayed pronounced cell-type-specific expression patterns across infected tissues (Fig. 3b). Despite these localized inductions, the global transcriptional output of the exocytosis pathway was broadly reduced in vascular tissues, including decreased expression of *GmSCD2*, most *GmEXO70* and *GmEXO84* isoforms, as well as *GmSNAP18 (Rhg1)* paralogs, *GmNSFs*, and several t-SNAREs. We hypothesize that this spatially restricted activation but overall repression of the exocytic pathway reflects a resistance mechanism mediated by the high-copy *Rhg1* QTL locus in PI 437654 (Patil *et al*., 2019a). The resistant allele of *GmSNAP18* has been shown to interact aberrantly with plasma-membrane t-SNAREs and NSF, impairing NSF-driven ATP hydrolysis and thereby preventing SNARE complex disassembly (Bayless *et al*., 2016; Dong, Zielinski & Hudson, 2020). Based on our data, we propose that in resistant roots carrying high-copy *Rhg1*, accumulation of *GmSNAP18* interferes with NSF-mediated SNARE recycling, leading to progressive failure of vesicle fusion cycles.

Together with our endocytosis results, these data suggest that SCN infection triggers a coordinated rewiring of host vesicle trafficking in which localized activation of exocytosis supports syncytium initiation, while simultaneous enhancement of endocytosis drives membrane turnover and cargo recycling required for feeding site expansion. However, in the Rhg1-mediated resistant background, accumulation of the SNAP18 protein disrupts NSF-dependent SNARE recycling, progressively collapsing exocytotic flux while CME-driven internalization continues. We therefore propose that an imbalance between exocytosis and endocytosis generates vesicle congestion and secretory stress within developing syncytial cells, ultimately destabilizing the feeding site and restricting nematode development.

### Endocycle suppression limits SCN feeding site formation

Endoreduplication represents a modified cell cycle in which DNA replication occurs without subsequent mitosis, leading to polyploidy and often accompanied by cell enlargement. Plant-parasitic nematodes have acquired the ability to induce remarkable changes in host cells during the formation of feeding sites (Reviewed: de Almeida Engler & Gheysen, 2013). Because SCN infection induces extensive cytological reprogramming in host roots (syncytium formation and nuclear hypertrophy) (de Almeida Engler & Gheysen, 2013; Breuer, Braidwood & Sugimoto, 2014; Sablowski & Gutierrez, 2022), we specifically examined endoreduplication-associated regulators to determine whether the resistant reaction constrains this reprogramming.

In our analysis, we observed differential expression of key regulators governing cell-cycle transitions (Fig. 4a, b). Transcripts corresponding to G1–S progression, including *GmE2FB* and *GmDEL1* (DP-E2F-Like1; (Heyman *et al*., 2017)), were highly enriched in xylem, indicating sustained mitotic competence. Genes driving S–G2 progression (*GmCDCs, GmORCs, GmMCMs*) were specifically expressed in cambium and developing syncytia, while G2–M regulators (*GmCYCA2*s, *GmCDKB2s*) showed predominant expression in xylem. Notably, *GmCDC20s*, a key activator of the anaphase-promoting complex (APC/C), displayed induction in pericycle cells, further supporting normal mitotic cycling rather than endocycle entry in this resistant genotype. In contrast, canonical endocycle-promoting genes (*GmSMR2* and *GmCCS52A*; Fig. 4) showed higher expression in control (SCN–) than in SCN+ xylem and phloem cells, suggesting a defense mechanism wherein the resistant plant actively suppresses endoreduplication, thereby preventing the nematode from inducing a large, metabolically active feeding site.

**Figure 4.**
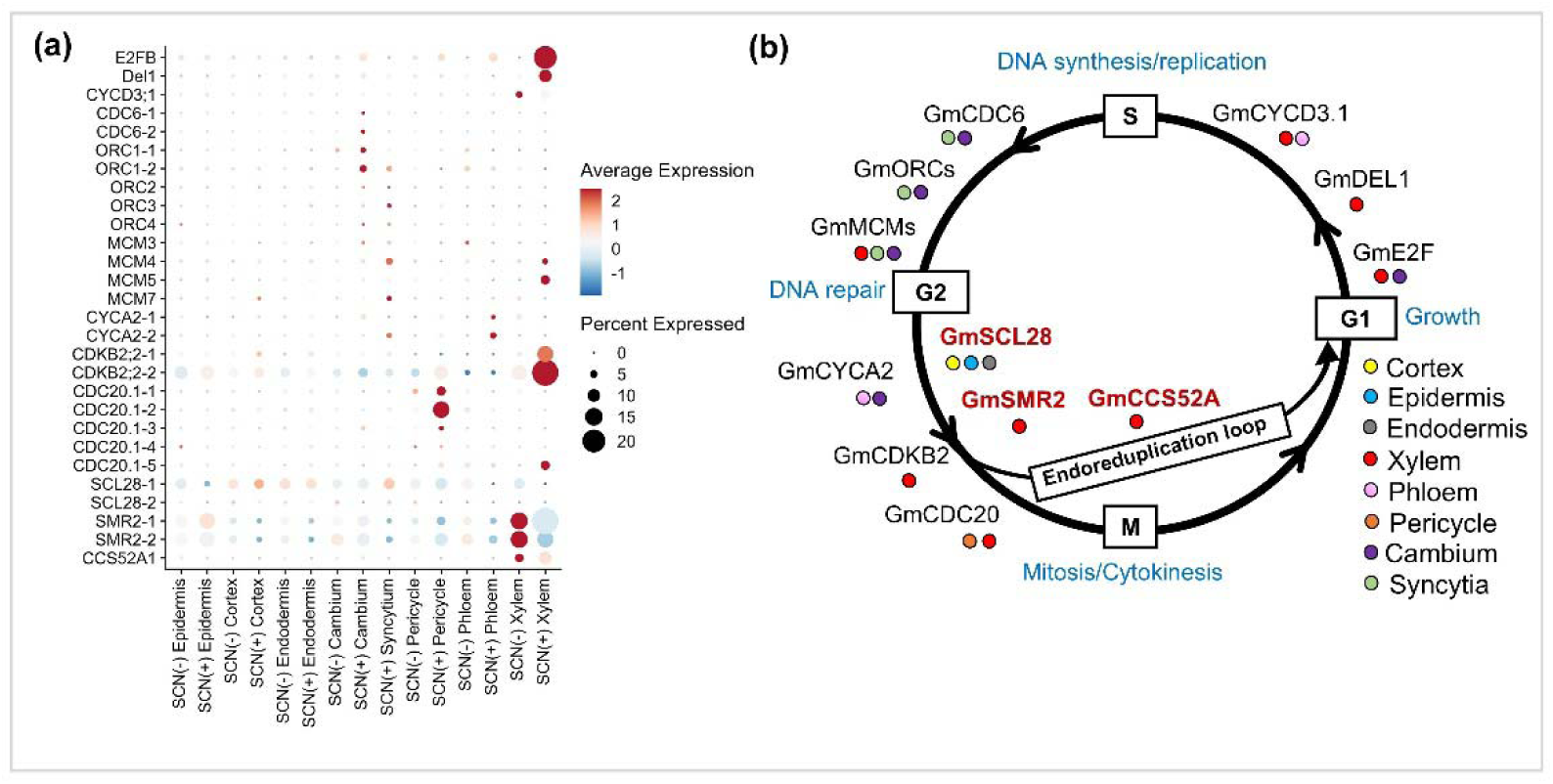
Cell-type-specific transcriptional responses of cell cycle and endoreduplication pathway. **(a)** A dot pot depicts transcript abundance of cell cycle and endoreduplication pathway genes after SCN (+) treatment. The colors indicate the relative expression. The size of the dots depicts the percentage of nuclei in each cluster that express the gene. The Red dots denote higher expression while blue dots exhibit lower expression. **(b)** Cartoon diagram illustrates cell cycle progression through G1-S-G2-M phases and the endoreduplication loop. Colored circles denote cell-type-specific gene expression patterns, as observed in SnRNA-seq dataset.

Building on this expression pattern, this suppression, coupled with the induction of the endocycle repressor *GmDEL1*, indicates an inhibition of the mitosis-to-endocycle transition in the resistant genotype PI437654. Consistent with their known roles, SCL28–SMR2 modules normally inhibit cyclin-dependent kinase activity to trigger endocycle onset (Yi *et al*., 2014; Kumar *et al*., 2015), while CCS52A activates APC/C to drive endoreduplication (Cebolla *et al*., 1999). Meanwhile, DEL1 acts antagonistically, repressing CCS52A2 to maintain mitotic cycling (Vlieghe *et al*., 2005). In Arabidopsis, loss of *CYCA2/CDKB2* or overexpression of *CCS52A* promotes endoreduplication, whereas *DEL1* overexpression restricts syncytium expansion and reduces nematode proliferation (de Almeida Engler et al., 2012; Imai et al., 2006; Endo et al., 2012). The strong induction of GmCYCA2, GmCDKB2, and GmCDC20 in our dataset aligns with a maintained mitotic state, mechanistically explaining the limited syncytial growth observed in resistant soybean roots. Together, these findings support a model wherein PI437654 enforces active mitotic cycling while repressing endocycle entry, thereby constraining syncytial polyploidization and nutrient sink formation required for SCN feeding site establishment which can be a central determinant of resistance.

### Targeted autophagy limits nematode-induced syncytium expansion

To dissect autophagy’s role during SCN infection at single-cell resolution, we analyzed expression of core autophagy genes in our snRNA-seq dataset (Fig. 5a, d; Table S4). Autophagy, a conserved intracellular degradation pathway, recycles damaged organelles and proteins to maintain cellular homeostasis under biotic stress (Liu & Bassham, 2012). The pathway progresses through sequential steps of initiation, nucleation, expansion, and maturation (Mizushima, 2007; Marshall & Vierstra, 2018). In our study, we observed cell-type-specific induction of autophagy initiation and nucleation components genes predominantly in cambium, xylem, and phloem cells (Fig. 5a, d). Genes mediating autophagosome expansion/maturation, including *GmATG8* (a–f) and the *GmATG12* conjugation system, were upregulated in phloem and cortical tissues, while closure-associated genes (*GmATG9*, *GmATG2*, *GmATG18*) showed the highest expression in xylem and phloem. Collectively, these patterns indicate that autophagy is selectively activated in vascular cell types upon SCN infection, suggesting a targeted role in mediating defense. To validate autophagy induction, we employed GFP-tagged GmATG8a, a well-established marker of autophagosomes (Yoshimoto *et al*., 2004; Marshall & Vierstra, 2018; Klionsky *et al*., 2021). Consistent with transcriptomic data, GFP–ATG8a puncta were elevated in vascular cells of infected roots compared to non-infected controls (Fig. 5b, c). Treatment with Concanamycin A (ConA), an inhibitor of vacuolar ATPases that blocks autophagosome degradation (Bassham, 2015), further increased puncta accumulation, confirming active autophagic flux in SCN-infected root cells. These results demonstrate that SCN infection triggers autophagosome formation and turnover, particularly in the vascular tissues critical for feeding site establishment.

**Figure 5.**
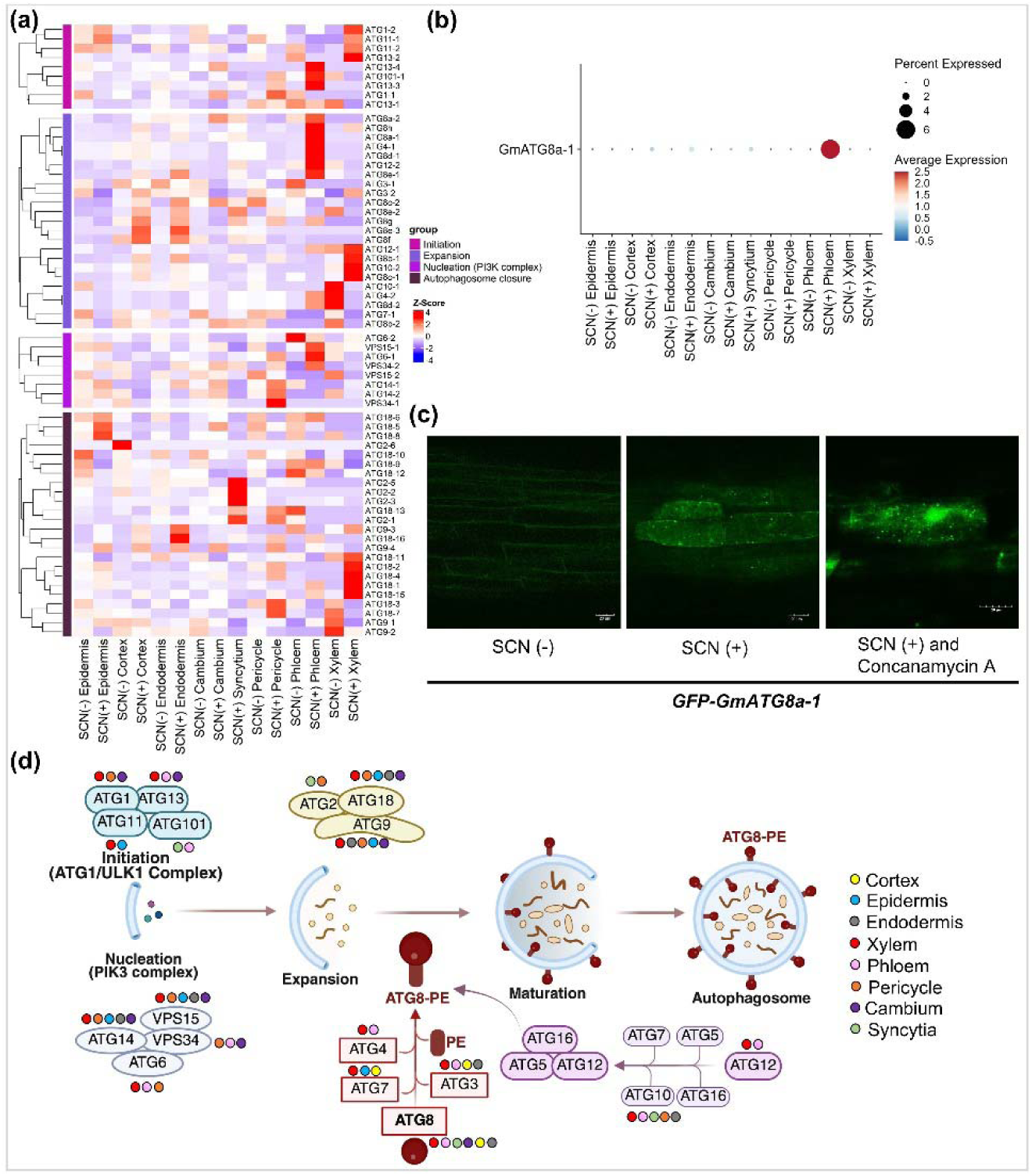
Cell-type-specific induction of autophagy during SCN infection. **(a)** Heatmap shows expression patterns of autophagy pathway genes and their regulatory components. Color scale represents relative expression levels; red indicates higher expression and purple indicate lower expression. **(b)** Dot plot shows expression of GmATG8a in SCN (-) and SCN (+) treated soybean roots in different cell types. **(c)** The autophagosome generation was observed after 5 days of SCN (+) infection in transgenic soybean hairy roots expressing *GFP-GmATG8a* fusion construct. The GFP-GmATG8a fluorescent puncta were detected in SCN (+) infected roots and roots treated concanamycin A. Bars, 20□μm. **(d)** Schematic model diagram exhibits the autophagy pathway and its important regulatory components across different cell types. Colored circles represent cell-type-specific gene expression patterns linked with the autophagy pathway.

Autophagy plays dual roles in plant-pathogen interactions (Hofius *et al*., 2017; Üstün, Hafren & Hofius, 2017). In plant-pathogen interactions, several pathogens secrete effectors to either hijack or stop autophagy to induce effective infection (Dagdas *et al*., 2016; Üstün, Hafren & Hofius, 2017). In the case of cyst nematodes, the effector Nematode Manipulator of Autophagy System 1 (NMAS1) harbors ATG8-interacting motif (AIM) that directly interacts with host plant ATG8 proteins to hijack their activity, indicating a vital role of the autophagy process (Chen *et al*., 2023). In tomato, autophagy enhances jasmonate-mediated resistance against root-knot nematodes, and its loss reduces host defense (Zou *et al*., 2023). Based on the induction of autophagy genes in PI437654 roots, particularly in feeding site-adjacent vascular cells, we hypothesized that, in this genotype, autophagy functions as a defense mechanism rather than being hijacked by the nematode. Elevated expression of ATG8 and other core autophagy components likely supports the selective degradation of nematode-induced effectors or damaged host components, limiting syncytium expansion and contributing to resistance. In summary, our data reveal that SCN infection triggers robust, cell-type-specific autophagy in soybean roots. In the resistant genotype PI437654, autophagy activation in vascular tissues appears to be a key component of defense, functioning to restrict feeding site development and support mitotic maintenance. This mechanistic link positions autophagy as an active regulator of nematode resistance and highlights its potential as a target for engineering enhanced SCN resilience in soybean.

### Spatial hormone networks govern nematode resistance

Phytohormone signaling constitutes a central process during plant–nematode interactions, where secreted nematode effectors actively rewire host hormonal circuitry to promote feeding-site while the host simultaneously initiates a defense response (Goverse & Bird, 2011; Kyndt, Fernandez & Gheysen, 2014; Gheysen & Mitchum, 2019). In our dataset of resistant reaction (PI437654), defense-responsive jasmonic acid (JA) biosynthesis genes were induced across multiple root cell types, including the epidermis, endodermis, cortex, and phloem, indicating a systemic activation of wound and defense-related response (**Figs 6a, S5**). In contrast, salicylic acid (SA) and ethylene biosynthesis genes showed their highest expression in phloem cells (**Figs 6a, S6**). This cell-type specific response aligns with established functional roles of these hormones in nematode resistance. JA acts as a positive regulator of nematode resistance, as evidenced by reduced infection following exogenous methyl-JA treatment and heightened susceptibility of JA-deficient mutants (Nahar *et al*., 2011; Kammerhofer *et al*., 2015; Gheysen & Mitchum, 2019). Similarly, SA mediates basal defense responses, with enhanced resistance following SA application and increased susceptibility in SA biosynthesis mutants (Wubben, Jin & Baum, 2008; Priya *et al*., 2011). The coordinated activation of JA across peripheral tissues together with SA and ethylene within vascular cells, therefore, suggests an active defense mechanism that simultaneously restricts penetration, feeding-site establishment, and systemic colonization in the resistant genotype.

**Figure 6.**
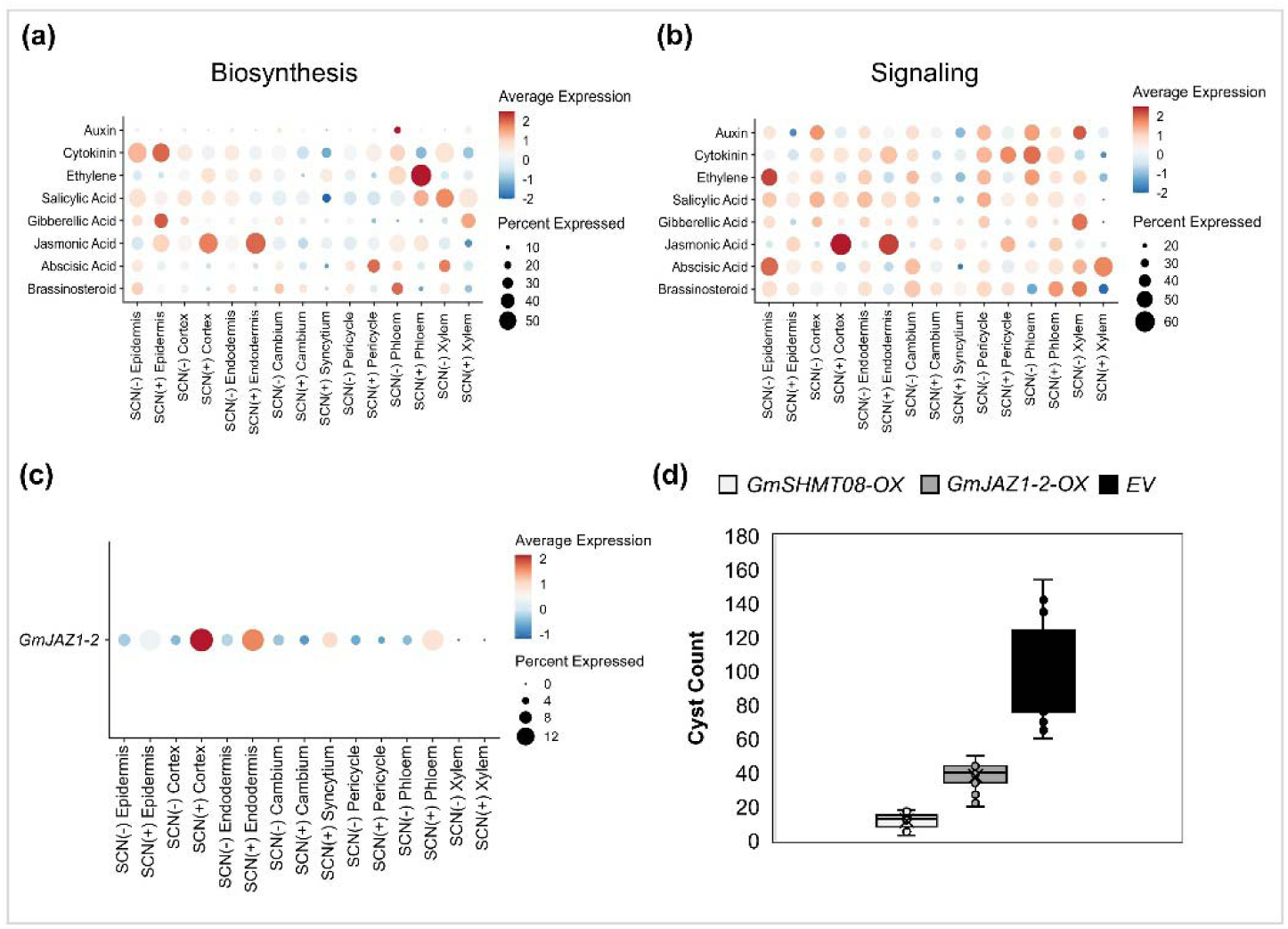
Cell-type-specific regulation of hormone biosynthesis and signaling and functional validation of GmJAZ1 during SCN infection. **(a)** Dot plot shows aggregated expression levels of phytohormone biosynthesis pathways, including auxin, cytokinin, ethylene, salicylic acid, gibberellic acid, jasmonic acid, abscisic acid, and brassinosteroid after SCN (-) and SCN (+) treatment. Each dot represents the combined expression of genes within a given hormone biosynthesis pathway. Dot size reflects the percentage of nuclei expressing genes from specific hormone pathway, while color indicates relative expression levels, red shows higher expression and blue shows lower expression. **(b)** Dot plot exhibits aggregated expression of phytohormone signaling pathways. Dot size indicates the percentage of nuclei expressing genes within each signaling hormone pathway, and color indicates relative expression levels, with red shows higher expression and blue indicate lower expression. **(c)** Dot plot shows expression of jasmonic acid singling master regulator *GmJAZ1* gene after SCN (-) and SCN (+) treatment. **(d)** SCN cysts were counted (Detection of SCN resistance) in composite plants having roots transformed with *A. rhizogenes* K599 carrying empty vector (EV; control), *GmJAZ1-OX* and *GmSHMT08 (Rhg4*)-OX constructs, respectively. Data are shown for the mean of twenty-six biological replicates ± SE for each construct. (Students t-test; * P < 0.05, ** P < 0.01).

In contrast to defense hormones, pathways associated with growth and developmental hormones associated with syncytium formation were selectively attenuated. Cytokinin (CK) and gibberellic acid (GA) biosynthesis genes exhibited induction in epidermal cells, likely reflecting early wound or penetration responses, but were suppressed in vascular tissues where feeding sites normally develop (**Fig. 6a**). Auxin and brassinosteroid (BR) biosynthesis genes were broadly downregulated across cell types following infection (Fig. 6a), indicating a global repression of growth-promoting signaling. This pattern is mechanistically significant because sedentary endoparasitic nematodes exploit these hormones to induce host cell cycle activation, hypertrophy, and syncytial expansion (Siddique *et al*., 2015; Yimer *et al*., 2018). Auxin accumulation at feeding sites and driven by redirected transport and local biosynthesis is essential for syncytium initiation, and disruption of auxin signaling impairs feeding-site development (Grunewald *et al*., 2009). Likewise, BR-mediated cell expansion contributes to feeding-site growth, and its suppression compromises nematode establishment (Nahar *et al*., 2013). Collectively, our data support a mechanistic model in which PI437654 resistance arises from a coordinated hormonal shift that favors defense over developmental reprogramming. Broad activation of JA, SA, and ethylene signaling likely creates a defensive environment for the nematode, while localized suppression of CK, GA, auxin, and BR deprives the nematode of the hormonal cues required to sustain feeding-site morphogenesis. This contrasting hormonal reprogramming effectively decouples infection from the tissue remodeling processes essential for nematode success, thereby restricting parasitism and conferring resistance.

### Functional validation identifies GmJAZ1 as cell-type–specific regulator of hormone crosstalk during SCN resistance

A comprehensive analysis revealed that among all phytohormone pathways, JA signaling exhibited the most spatially coherent activation upon SCN infection, particularly in cortex and endodermal layers that constitute the primary invasion route of the nematode (**Figs 6b, S5**). Within this pathway, members of the *GmJAZ* (*Jasmonate ZIM-domain*) family emerged as the most consistently and highly induced regulators across cortex, endodermis, and phloem cells **(Fig. S5)**. JAZ proteins are central repressors of JA signaling that directly control the activity of key transcription factors such as MYC2, thereby functioning as molecular nodes that integrate JA signaling with other hormone pathways (Jing *et al*., 2019). We prioritized *GmJAZ1-2* for functional validation because (i) it showed robust induction across multiple infection-relevant cell types rather than a single compartment, (ii) it occupies an upstream regulatory position capable of modulating entire downstream defense programs, and (iii) JAZ proteins are well-established mediators of JA–SA antagonistic crosstalk, a process central to defense against biotrophic pathogens. In contrast, downstream biosynthetic or response genes typically exert context-dependent or redundant effects, making them less suitable for mechanistic interrogation of pathway control. JA and SA constitute two major immune signaling hubs that often act antagonistically to tailor defense responses to specific pathogen lifestyles (Glazebrook, 2005; Pieterse *et al*., 2012). SA-dependent pathways generally confer resistance to biotrophic pathogens, whereas JA/ethylene pathways are associated with defense against necrotrophs and herbivorous insects. Mechanistically, activation of JA signaling through MYC2 and ERF transcription factors suppresses SA-responsive genes (e.g., PR1, ICS1), while SA accumulation reciprocally represses JA-responsive genes (Spoel *et al*., 2003). Such reciprocal inhibition generates highly specialized defense outputs but can also be manipulated by pathogens to facilitate infection. Given that SCN is an obligate biotroph, modulation of this JA–SA balance is likely a critical determinant of host outcome.

To test whether JAZ-mediated regulation of this hormonal node influences SCN infection, we functionally characterized *GmJAZ1-2* using composite transgenic soybean roots (Devkar *et al*., 2025a). *GmJAZ1-2* expression was strongly induced in cortex, endodermis, and phloem tissues following infection **(Fig. 6c)**. Overexpression of *GmJAZ1-2* in the susceptible cultivar Williams 82 (W82) conferred a pronounced resistance phenotype, exhibited by a significant reduction in cyst formation relative to empty-vector controls **(Fig. 6d)**. This result demonstrates that elevating a single upstream regulator is sufficient to shift the host environment toward an anti-nematode state, highlighting the regulatory leverage of JAZ proteins within the hormone network. To further interrogate gene function, we generated CRISPR/Cas9 knockouts of *GmJAZ1-2* using dual gRNAs targeting exon 1 (Fig. S7a). An 88-bp deletion was recovered in GFP-positive composite roots (Fig. S7b), confirming efficient mutagenesis. However, knockout of *GmJAZ1-2* in the resistant genotype PI437654 did not significantly alter cyst numbers compared with controls (Fig. S7c). The soybean genome encodes multiple JAZ paralogs (Table S4), and extensive functional redundancy among JAZ proteins has been documented in plants (Chini *et al*., 2016). Thus, loss of a single family member is likely buffered by compensatory activity of other GmJAZ proteins, whereas overexpression can overcome this redundancy by quantitatively shifting repression of JA signaling. Mechanistically, the resistance phenotype observed upon *GmJAZ1-2* overexpression is consistent with a model in which enhanced JAZ levels attenuate JA signaling, thereby relieving suppression of SA-mediated defenses that are more effective against biotrophic pathogens (Jing *et al*., 2019). Supporting this interpretation, JAZ overexpression in soybean has been shown to suppress JA–MYC2 activity and promote SA-dependent responses. For example, overexpression of *TaJAZ1* in wheat increased resistance to the biotrophic fungus causing powdery mildew and elevated expression of SA-responsive genes such as *TaPR1/2* (Jing *et al*., 2019). Similarly, JAZ-mediated repression of JA signaling can activate SA-dependent immunity through hormonal antagonism (Spoel *et al*., 2003; Gimenez-Ibanez & Solano, 2013; Nomoto *et al*., 2021). In summary, our results identify GmJAZ1 as a strategic regulatory node controlling hormone crosstalk during SCN infection. We propose that in soybean, infection-induced upregulation of GmJAZ1 dampens JA signaling in key invasion tissues, thereby enabling activation of SA-mediated defenses that restrict biotrophic nematode establishment. This mechanism provides a coherent explanation for the strong resistance phenotype observed upon overexpression and underscores the importance of regulatory, rather than merely biosynthetic, components of hormone pathways in determining host outcome.

## Conclusion

This study provides a high-quality, cell-type–resolved atlas of soybean root responses during SCN infection in a highly resistant genotype, delivering a mechanistic view of how host cellular programs are reconfigured to block parasitism. We generated a high-resolution dataset capturing transcriptional states of all major root tissues, including rare and previously inaccessible syncytial cells **(Fig. 2)**. This resource reveals that SCN resistance in PI437654 is not mediated by a localized defense reaction but by a coordinated, multi-cell reprogramming of cellular functions spanning invasion layers, vascular tissues, and feeding site–associated cells. A central advance of this work is the identification of vascular cambium as the primary cellular origin of SCN-induced syncytia, resolving a long-standing question in nematology. This model suggests that nematode-induced remodeling extends beyond the feeding cell itself and involves surrounding cambial cells that either facilitate expansion or mount counter-defense responses.

Mechanistically, resistance in PI437654 emerges from a coordinated disruption of the cellular processes required for feeding site establishment. This study revealed that SCN infection triggers extensive remodeling of vesicle trafficking pathways, with activation of clathrin-mediated endocytosis and endosomal transport in vascular and syncytial cells **(Fig. 3)**. Exocytosis is globally attenuated in the resistant background, consistent with interference by the high-copy *Rhg1* locus and accumulation of SNAP18 proteins that impair SNARE recycling (Bayless *et al*., 2016). We propose that this imbalance between endocytosis and exocytosis generates secretory stress and vesicle congestion, destabilizing syncytial function and limiting nematode development. This supports a mechanistic explanation for *Rhg1*-mediated resistance at the cellular level.

In parallel, resistant roots actively block nematode-induced developmental reprogramming. While successful infection requires endoreduplication and hypertrophic growth, PI437654 maintains mitotic cycling and suppresses the mitosis-to-endocycle transition, preventing polyploidization and formation of a strong nutrient sink (Fig. 4). Concurrent activation of autophagy in vascular tissues likely removes nematode effectors or damaged components, further restricting syncytial expansion, indicating resistance is enforced through active cellular homeostasis rather than passive tolerance (Fig. 5). We also uncover a spatially organized hormone defense network that favors immunity over the growth. JA, SA, and ethylene pathways were activated in a tissue-specific manner. Functional validation identifies *GmJAZ1* as a central regulator of JA–SA crosstalk, where its overexpression confers resistance, demonstrating that upstream hormone regulators can reprogram infection outcomes.

Together, these data support a unified model in which SCN resistance disrupts three essential requirements for parasitism: permissive host cell identity, formation of a metabolically active feeding site, and sustained nutrient acquisition. Accordingly, PI437654 deploys a multilayered defense strategy targeting multiple stages of the nematode life cycle. Beyond mechanistic insights, this work establishes a high-resolution single-cell resource for soybean roots under biotic stress, capturing rare cell states and regulatory networks inaccessible to bulk approaches. Overall, our study advances the field from descriptive observations to a systems-level framework explaining how resistant hosts actively block nematode parasitism and highlights actionable targets for engineering durable SCN resistance.

## Materials and Methods

### Plant materials

The snRNA-seq experiments demonstrated in this study were performed in the broad-spectrum cyst nematode-resistant soybean accession PI 437654. These plants were grown in a controlled greenhouse and maintained at ∼26°C temperature, 60-70% relative humidity, a photoperiod of 16 hours light and 8 hours dark. Soybean cyst nematode HG type 0 (race 3) has been multiplied on “Hutcheson” and “Lee 74” nematode susceptible lines.

### Soybean cyst nematode treatment and phenotyping

SCN Phenotyping was conducted in a controlled greenhouse environment at the University of Missouri, Columbia, using a standardized bioassay (Niblack *et al*., 2009), with modifications as described earlier (Chhapekar *et al*., 2025). Briefly, seeds of PI 437654 and indicator lines were germinated in paper pouches for 4 days and transplanted into PVC micro pots containing steam-pasteurized sandy soil. Indicator lines include PI 88788, PI 437654, Peking, PI 209332, PI 90763, PI 89772, PI 548316, and Pickett with susceptible controls ‘Hutcheson’ and ‘Lee 74’ were included in this experiment. Micropots were arranged in plastic containers and maintained in water baths at 27°C. Seedlings were organized in a randomized complete block design, with 10 plants per genotype. Two days after transplanting, each plant was inoculated with approximately 2,000 SCN eggs from HG type 0 (race 3).

At 30 days post-inoculation, root systems were thoroughly washed, and cysts were gently collected using a 250 µm sieve and quantified with fluorescence-based imaging system. Resistance responses were quantified using the female index (FI), calculated as the percentage ratio of the mean number of female cysts on a test genotype relative to the susceptible control. Each genotype was evaluated with more than ten biological replicates, and experiments were conducted at least three independent times across different seasons. Final FI values represent the mean across all experiments. Soybean accessions were classified as resistant (FI < 10; R), moderately resistant (FI = 11–30, MR), moderately susceptible (FI = 31–60, MS), or susceptible (FI > 60, S) (Schmitt and Shannon 1992).

### Confirmation of soybean cyst nematode penetration through staining soybean roots

Nematode penetration and early infection were examined in PI 437654 roots to confirm successful SCN infection. Seedlings were inoculated with SCN HG type 0 as described above. At 5 days after inoculation (dai), root systems were carefully removed from the soil, and five biological replicates containing SCN infected root system were collected per accession. Fresh roots were processed using an acid fuchsin staining method with minor modifications (Bybd Jr, Kirkpatrick & Barker, 1983). Roots were washed thoroughly with tap water, treated with 5.25% sodium hypochlorite (NaOCl) for 4 min with gentle agitation to remove residual debris, and rinsed again with water. Samples were then briefly boiled for about 30 seconds in 37% acid fuchsin staining solution to visualize nematodes, followed by rinsing and destaining in glycerol. Stained roots were examined under a dissecting microscope, and juvenile nematodes (J2/J3) were identified based on morphological characteristics.

### Nuclei isolation, SnRNA-seq library preparation, and sequencing

Nuclei isolation was performed from SCN(+) and SCN(-) treated soybean root tissues as previously described (Devkar *et al*., 2025a). Freshly collected root samples were immediately chopped with a razor blade in a nuclei isolation buffer and incubated briefly (2-3 min) on a shaker at 60 rpm, followed by passing through 100 µm, 70 µm, 30 µm, and 10 µm cell strainers. The nuclei pellet was resuspended in wash buffer and spun at 1000g for 2 minutes. The nuclei were stained with propidium iodide and counted with the Luna-FL cell counter (Logos Biosystems). The nuclei were then loaded onto the chip as per the manufacturer’s recommendations to target 10,000 nuclei recovered (10X Genomics, Pleasanton, CA, USA). Library construction for Illumina sequencing was performed with the Chromium™ Single Cell 3’ Library & Gel Bead Kit v3.1 protocol (10X Genomics). The single-indexed sequencing of paired-end libraries were carried out on an Illumina™ NovaSeq 6000 platform according to the 10X Genomics manual.

### SnRNA-seq data pre-processing and clustering

Sequencing data was processed using kallisto bustools (Melsted *et al*., 2021) to obtain cell feature counts for analysis. The data was aligned to the Glycine max v4 genome. Count data was further analyzed within the R package Seurat v4 (Hao *et al*., 2021). Chloroplast and mitochondrial reads were removed, and nuclei in the 95th percentile for gene count and 90^th^-99^th^ percentile for UMI were selected for further pre-processing. Then, doublets were removed with scDblFinder (Germain *et al*., 2022), and background RNA was removed with SoupX (Young & Behjati, 2020). A median UMI of 295 and 362, and a median number of genes of 513 and 592 were obtained for SCN(+) and SCN (-), respectively. The data was then normalized by SCTransform, and the samples were integrated using Harmony (Korsunsky *et al*., 2019) using PCs 1 to 30 and group.by.vars = “treatment” to reduce batch effect and identify conserved cell types. Dimension reduction of the integrated data was performed using harmony reduction, and the UMAP plots were generated using dims 1 to 30. Monocle3 clustering was applied to cluster cells and identify the potential cell types (Cao *et al*., 2019).

### Cell type annotation

Annotation of cell types was performed by utilizing the published soybean single-cell markers from (Liu et al., 2023; Sun et al., 2023; Cervantes-Pérez et al., 2024; Zhang et al., 2025). Expression of the markers was analyzed using the AddModuleScore function from Seurat and observed using the dotplot and featureplot functions. SCN induced marker genes were obtained using the FindAllMarkers function with min.pct = 0.2, logfc.threshold = 0.2, min.diff.pct = 0.2. GO term enrichment was performed using the topGO package in R (Alexa A, 2025).

### Vasculature Analysis

Vasculature cells were subset for further analysis and re-clustered using PCs 1 to 10. Cambium clusters were then subset further due to high expression of syncytium marker genes. Re-clustering of cambium genes was performed using Monocle3’s reduce_dimension command and trajectory analysis were performed using learn_graph (Cao *et al*., 2019). Trajectory DEGS were clustered using find_gene_modules and top DEGS were visualized using complexheatmap (Gu, Eils & Schlesner, 2016).

### Constructs and cloning

To functionally validate GmJAZ1 in soybean against SCN pest, we generated overexpression of *GmJAZ1* by cloning *GmJAZ1* gene (Synthesized by Twist Bioscience, USA) under the control of constitutive promoter *CmYLCV (GmJAZ1-OX)* using gateway cloning. The fluorescence marker GFP was expressed under *CmYLCV* promoter (pMod_C3003) for detection of transgenic roots, these cassettes were assembled via Golden Gate in binary vector pTrans230d. We used SHMT08-overexspression (Rhg4 source of resistance) construct (Lakhssassi *et al*., 2025) as positive control for SCN resistance. Later we generated composite transgenic hairy roots using *Agrobacterium rhizogenes K599.* For editing *GmJAZ1* by CRISPR/Cas9, Golden Gate construct modules namely A0521 (AtCas9), B2103b (cloning dual guide RNAs), C3003 (GFP marker) and pTRANS230d (binary vector) were used (Čermák *et al*., 2017). The guide RNAs (gRNAs) were designed using CRISPR-P tool (Liu *et al*., 2017). High-ranking gRNAs were selected with no possible off-targets (Liu *et al*., 2017). Two gRNAs were designed with a 88 bp distance to induce larger deletions. Genotyping was performed as explained (Devkar *et al*., 2025a; Devkar *et al*., 2025b). For autophagy detection and validation of autophagosome, we synthesized eGFP-GmATG8a gene block from Twist Bioscience, USA. Later, we cloned *eGFP-GmATG8a* in entry clone pd221 vector using BP clonase enzyme via Gateway cloning. By using LR clonase and gateway recombination method, *eGFP-GmATG8a* moved to destination vector to express under GmUbi promoter (pMod_A215). To detect transgenic hairy roots, we used module C3003-RFP vector having RFP expressed under CmYLCV promoter. Finally, by using Golden Gate cloning all modules including A215, B000 and C3003-RFP were assembled in binary vector pTrans230d as described (Čermák *et al*., 2017). All plasmids used in this study were sequence confirmed by sequencing whole plasmids by Oxford Nanopore Technology from Plasmidsaurus, Eugene, USA.

### Generation of transgenic hairy, composite soybean roots and SCN bioassay

Transgenic soybean hairy roots were generated using *Agrobacterium rhizogenes* (*strain, K599*)-transformation method as explained (Devkar *et al*., 2025b). Composite soybean plants with transgenic roots were generated using *A. rhizogenes* (*strain, K599*)-facilitated transformation as per the established protocol (Fan *et al*., 2020). Briefly, Composite hairy roots were generated in soybean cultivar Williams 82 via Agrobacterium rhizogenes strain K599, injected into the hypocotyl region below the cotyledons. For each construct, composite roots were obtained from at least 50 independent plants and grown in vermiculite under high-humidity conditions for 2 weeks. Plants were fertilized weekly (NPK 20–20–20), and those bearing GFP-positive roots (2–3 inches in length) were transferred to sandy soil for SCN screening. SCN bioassays were performed using HG type 0 (race 3), the most prevalent race in U.S. soybean fields, as per the method described above. At 30 days post-inoculation, cysts were counted under a stereomicroscope. Each experiment included about 20 to 25 independent transgenic root lines per construct and was repeated three times independently. Data was analyzed by analysis of variance (ANOVA).

#### Autophagosome detection

Confocal microscopy-based identification of autophagosomes in soybean hairy roots was performed as described previously (Thompson *et al*., 2005). Briefly, soybean hairy roots over-expressing *GFP-GmATG8a* were infected with SCN as described above in SCN infection assay section, then control and SCN infected roots were cut into small sections, and autophagosomes were observed with an Olympus confocal microscope with excitation at 488□nm and emission at 493 to 558□nm. Concanamycin A (ConcA) of 1□μM was applied to roots for 3□h, before confocal microscopy.

## Supplemental Figure Legends

**Supplemental Fig. S1.** UMAP exhibits diverse cell types upon SCN infection in resistant genetic source PI437654 roots.

**Supplemental Fig. S2.** UMAP plot exhibits expression of cell-type-specific representative marker genes. The color depicts relative gene expression (dark purple = higher expression, Orange = medium expression, Yellow = lower expression).

**Supplemental Fig. S3.** Gene Ontology (GO) enrichment analysis for the marker genes within each cluster of SCN (-) and SCN (+) infected root samples. X-axis exhibits GO terms, while Y-axis shows different cell types. Size of circles represent number of genes. Whereas blue to red color denotes GO enrichment.

**Supplemental Fig. S4.** UMAP (Upper panel) and dot plot (lower panel) shows expression of syncytia specific expressed genes captured by laser capture microdissection (LCM) from Klink et al (2005; 2007 and 2010) in different cell types upon SCN infection. The colors exhibit the relative gene expression. (blue = lower expression, dark Red = higher expression). The size of the dots represents percentage of cells genes expressed.

**Supplemental Fig. S5.** Dot plot shows expression of Jasmonic acid (JA) biosynthesis and signaling genes in SCN (-) and SCN (+) treated samples in different cell types. The colors denote the relative gene expression. (blue = lower expression, dark Red = higher expression). The size of the dots represents percentage of cells genes expressed.

**Supplemental Fig. S6.** Dot plot represents expression of Salicylic acid (SA) biosynthesis and signaling genes in SCN (-) and SCN (+) treated samples in various cell types. The colors show the relative gene expression. (blue = lower expression, dark Red = higher expression). The size of the dots exhibits percentage of cells genes expressed.

**Supplemental Fig. S7.** Functional characterization of GmJAZ1a CRISPR/Cas9 knockout during SCN infection. (a) The upper panel cartoon image displays CRISPR/Cas9 dual guide RNA (gRNA) target location on *GmJAZ1* gene. Upended red colored triangles denote gRNAs, whereas gray boxes depict exons and middle thick line joints are introns; bottom panel cartoon depicts CRISPR/Cas9 construct design. (b) CRISPR/Cas9 genome editing of soybean *GmJAZ1-2* (*CR-GmJAZ1-2*) in transgenic roots of composite soybean plants. NGS-Amplicon sequencing (205,248 reads) results analyzed by CRISPResso2 tool exhibits gene editing results by genotyping of amplicon from transgenic roots of composite plants transformed with *A. rhizogenes K599* carrying *CR-GmJaz1-2* construct. (c) SCN cysts were counted (Detection of SCN resistance) in composite plants having roots transformed with *A. rhizogenes* K599 carrying empty vector (EV; control) and CR-*GmJAZ1-2* constructs. Data are shown for the mean of Nineteen biological replicates ± SE for each construct.

## Data availability statement

All soybean gene IDs related to the genes evaluated in this study are specified in Supplemental Table S4. The single-nucleus RNA-sequencing data produced will be provided upon request to the corresponding author.

## Supplemental Information

Supplemental Figs (S1 – S7)

Supplemental Tables (S1-S4)

## Author Contributions

VD: experimental design, formal analysis, conceptualization, gene-editing, genotyping, confocal microscopy, data prediction and writing first draft; LDA: snRNA-seq analysis, formal analysis, data interpretation, methodology and contributed in writing first draft; SSC: methodology, composite hairy roots, SCN infection and phenotyping and manuscript editing; JL: SCN infection, confocal microscopy; MF: confocal microscopy; LY-V: snRNA-seq library preparation; LHE: supervision and editing manuscript; FLG: supervision and editing manuscript; HN: supervision and editing manuscript GBP: Conceptualization, experimental design, formal analysis, funding acquisition, supervision and writing and editing manuscript.

## Acknowledgements

GBP is grateful to the United Soybean Board, United States Department of Agriculture (USDA) and the State of Texas’ Governor’s University Research (GURI) for the research funding.

## Declaration of Interests

Authors declare no competing interests.

## Notes

### Competing Interest Statement

The authors have declared no competing interest.

