## Supplementary Figures for "Cell-type-resolved transcriptional reprogramming in resistant soybean roots reveals cambial activation and early syncytium initiation upon nematode infection"

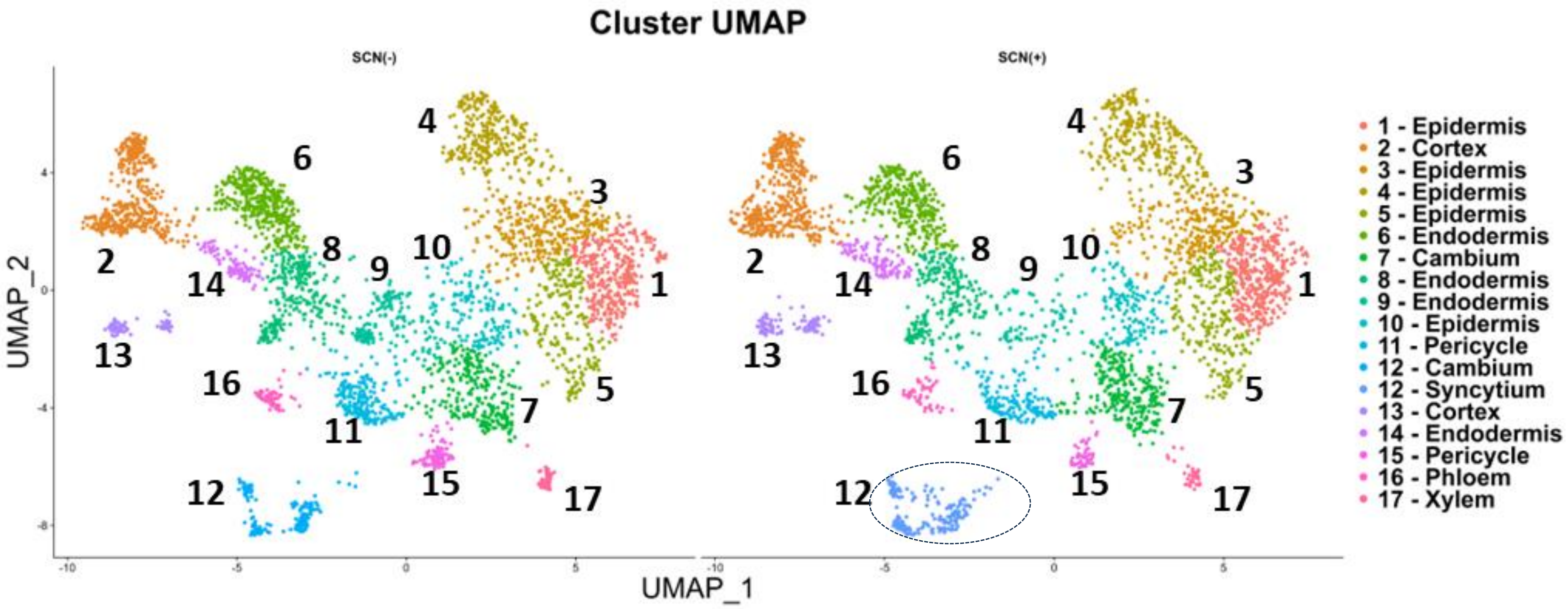


**Supplemental Fig. S1. UMAP exhibits diverse cell types upon SCN infection in resistant genetic source PI437654 roots.**

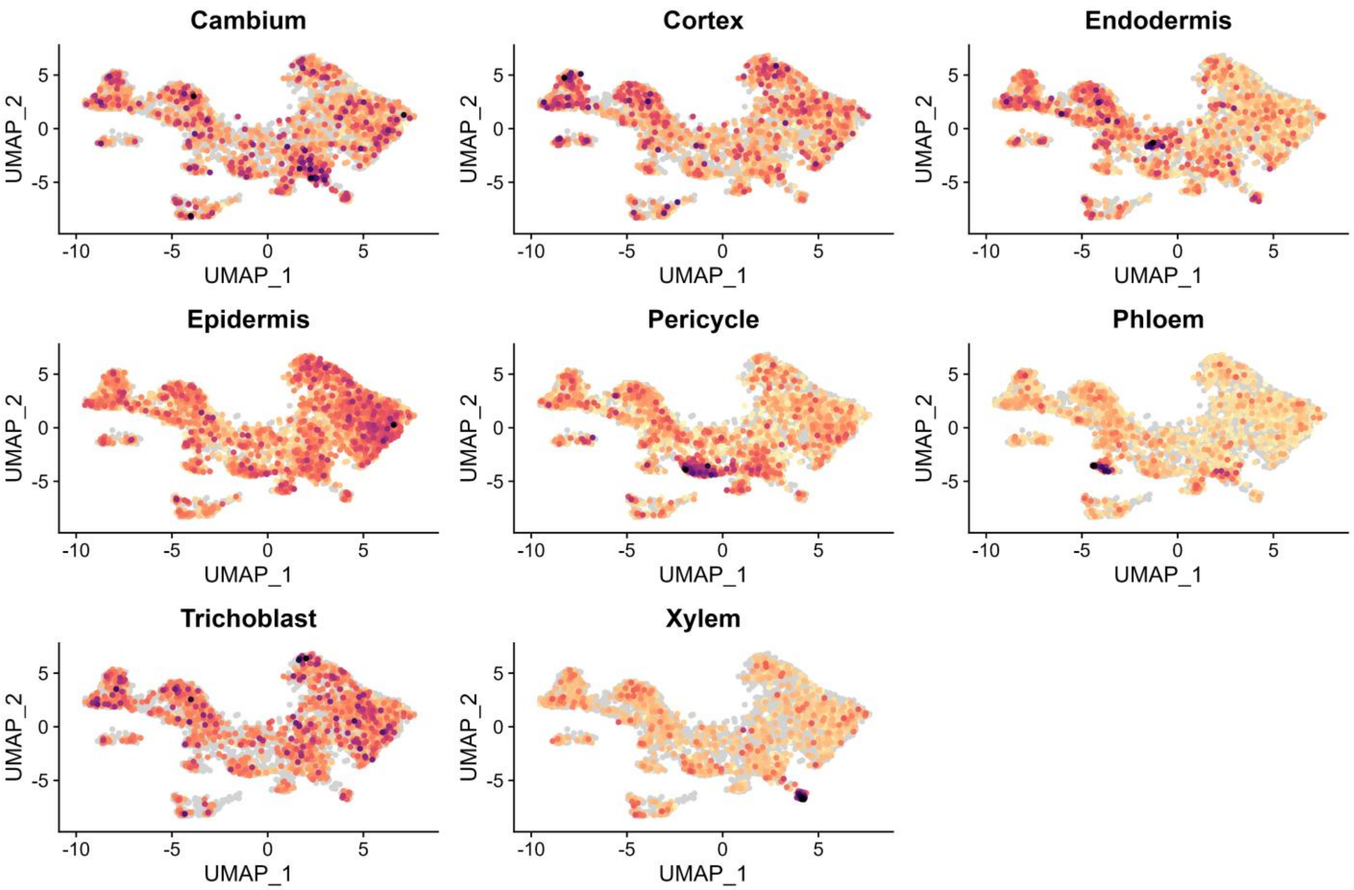


**Supplemental Fig. S2. UMAP plot exhibits expression of cell-type-specific representative marker genes.** The color depicts relative gene expression (dark purple = higher expression, Orange = medium expression, Yellow = lower expression).

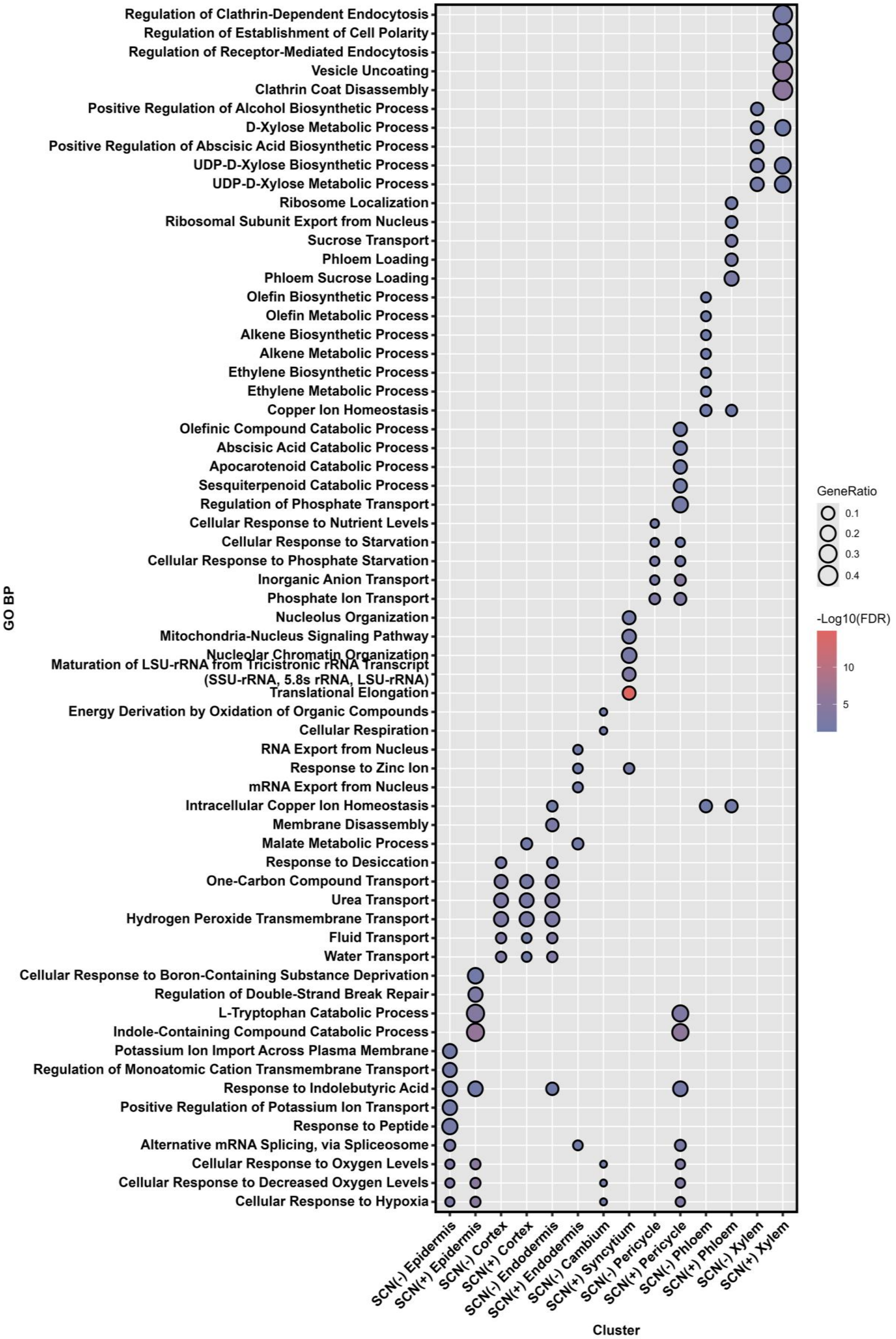


**Supplemental Fig. S3. Gene Ontology (GO) enrichment analysis for the marker genes within each cluster of SCN (-) and SCN (+) infected root samples. X-axis exhibits GO terms, while Y-axis shows different cell types. Size of circles represent number of genes. Whereas blue to red color denotes GO enrichment.**

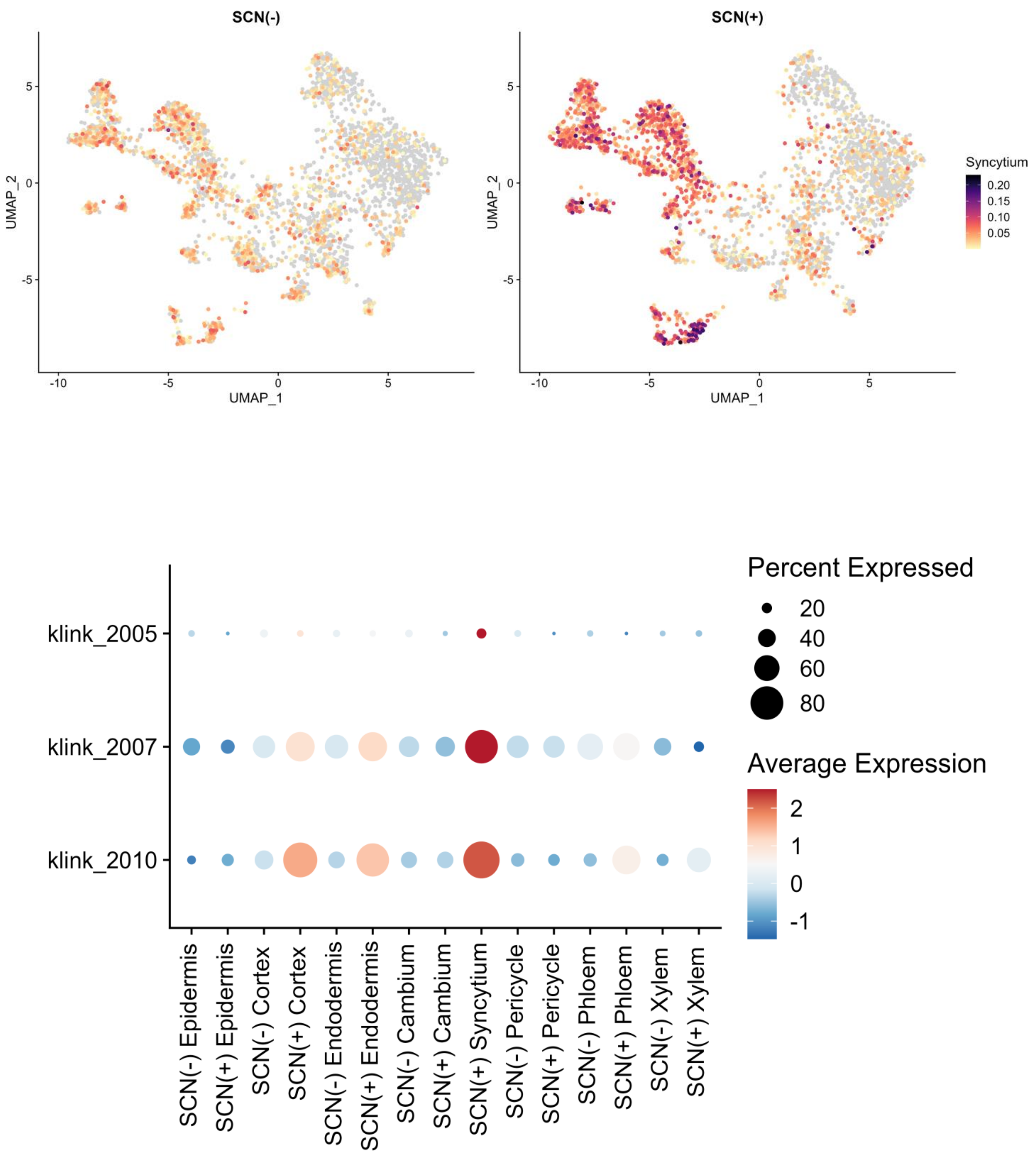


**Supplemental Fig. S4. UMAP (Upper panel) and dot plot (lower panel) shows expression of syncytia specific expressed genes captured by laser capture microdissection (LCM) from Klink et al (2005; 2007 and 2010) in different cell types upon SCN infection. The colors exhibit the relative gene expression. (blue = lower expression, dark Red = higher expression). The size of the dots represent percentage of cells genes expressed.**

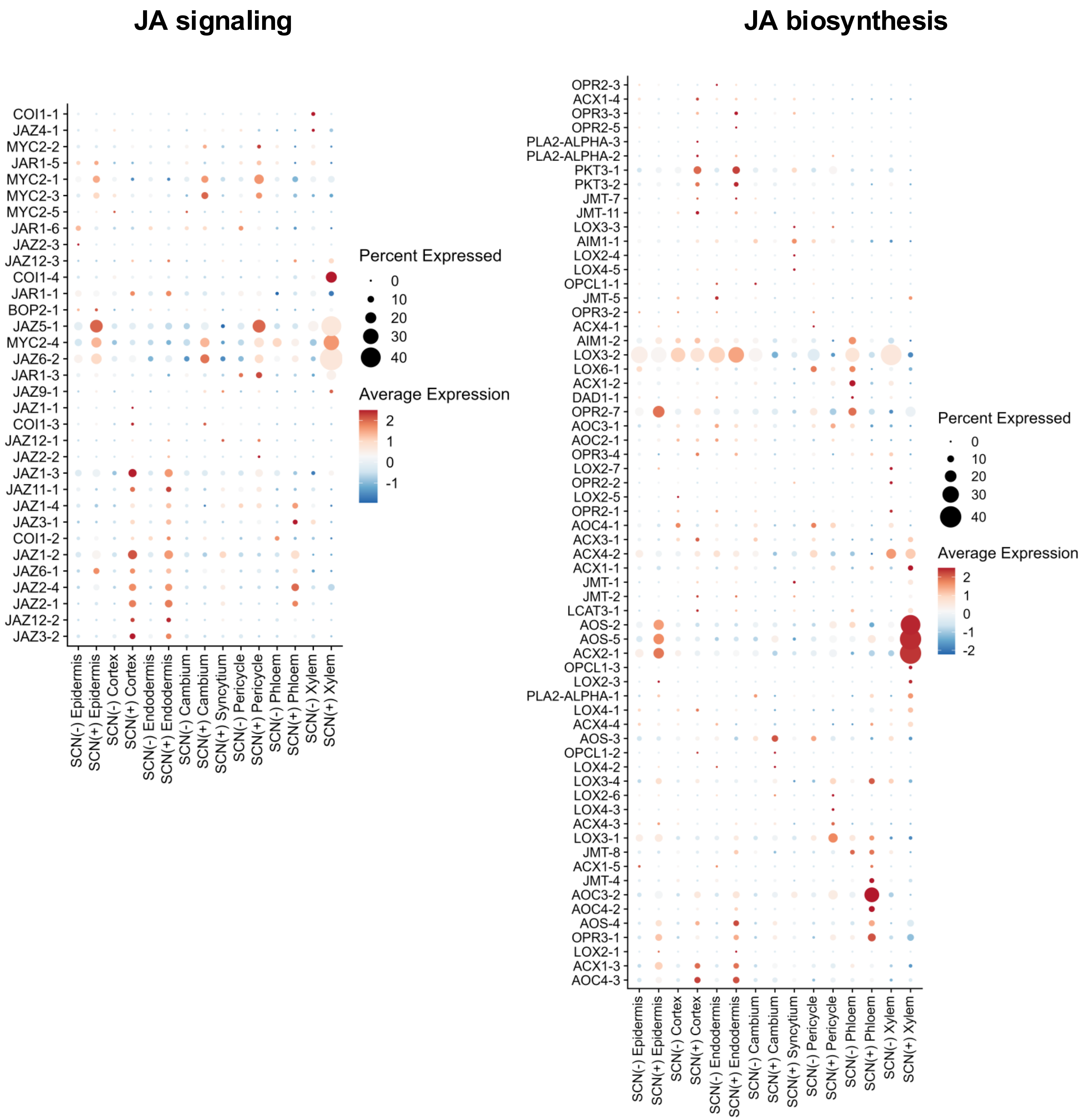


**Supplemental Fig. S5. Dot plot shows expression of Jasmonic acid (JA) biosynthesis and signaling genes in SCN (-) and SCN (+) treated samples in different cell types. The colors denotes the relative gene expression. (blue = lower expression, dark Red = higher expression). The size of the dots represent percentage of cells genes expressed.**

### SA signaling

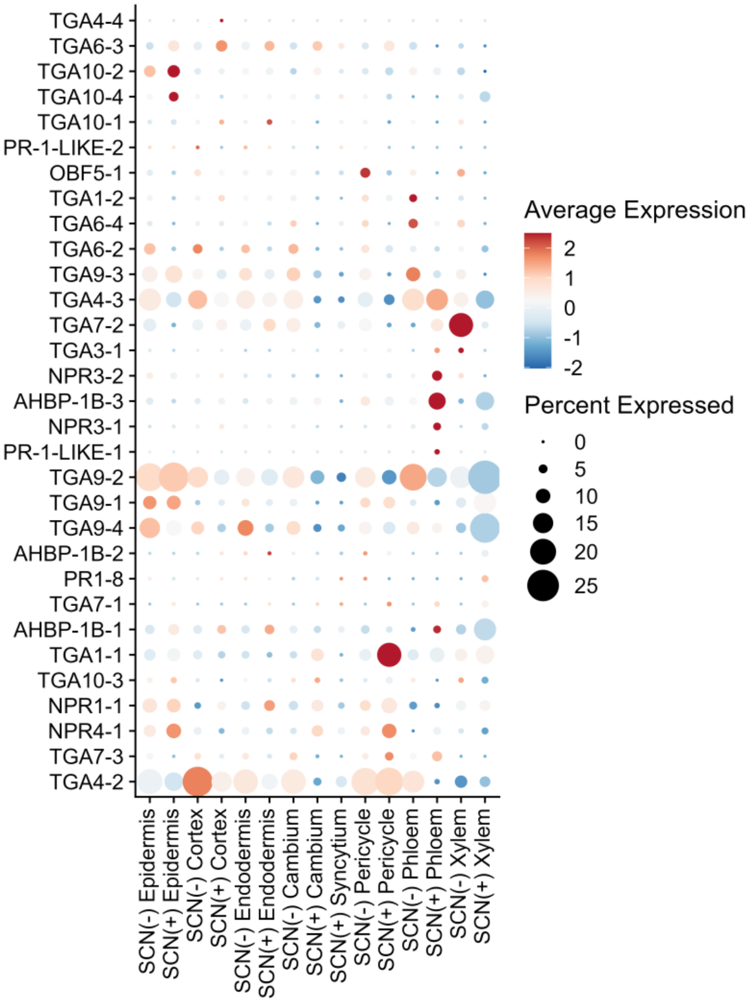


### SA biosynthesis

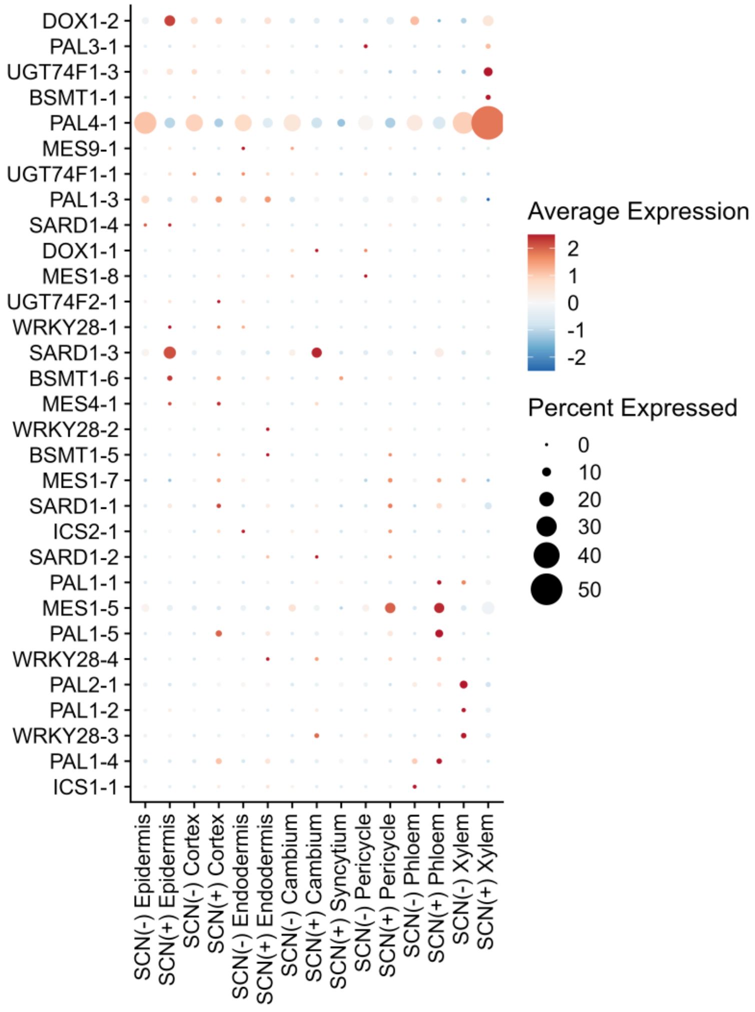


**Supplemental Fig. S6. Dot plot represents expression of Salicylic acid (SA) biosynthesis and signaling genes in SCN (-) and SCN (+) treated samples in various cell types. The colors shows the relative gene expression. (blue = lower expression, dark Red = higher expression). The size of the dots exhibits percentage of cells genes expressed.**

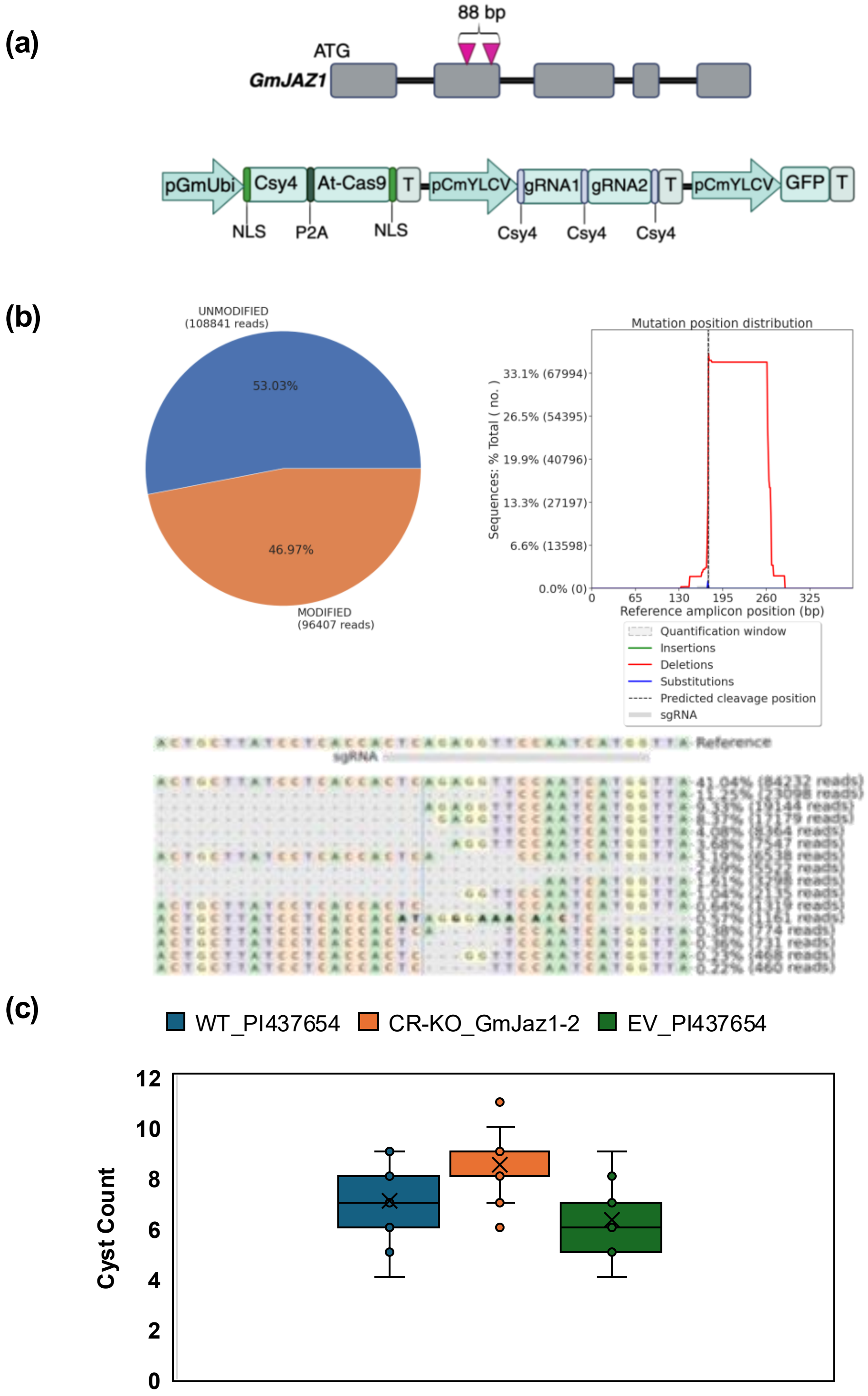


**Supplemental Fig. S7 Functional characterization of GmJAZ1a CRISPR/Cas9 knockout during SCN infection.**

(a) The upper panel cartoon image displays CRISPR/Cas9 dual guide RNA (gRNA) target location on *GmJAZ1* gene. Upended red colored triangles denote gRNAs, whereas gray boxes depict exons and middle thick line joints are introns; bottom panel cartoon depicts CRISPR/Cas9 construct design. (b) CRISPR/Cas9 genome editing of soybean *GmJAZ1-2* (*CR-GmJAZ1-2*) in transgenic roots of composite soybean plants. NGS-Amplicon sequencing ( 205,248 reads) results analyzed by CRISPResso2 tool exhibits gene editing results by genotyping of amplicon from transgenic roots of composite plants transformed with *A. rhizogenes K599* carrying *CR-GmJaz1-2* construct. (c) SCN cysts were counted (Detection of SCN resistance) in composite plants having roots transformed with *A. rhizogenes* K599 carrying empty vector (EV; control) and CR-*GmJAZ1-2* constructs. Data are shown for the mean of Nineteen biological replicates ± SE for each construct.
